# Microglia monitor and protect neuronal function via specialized somatic purinergic junctions

**DOI:** 10.1101/606079

**Authors:** Csaba Cserép, Balázs Pósfai, Barbara Orsolits, Gábor Molnár, Steffanie Heindl, Nikolett Lénárt, Rebeka Fekete, Zsófia I. László, Zsolt Lele, Anett D. Schwarcz, Katinka Ujvári, László Csiba, Tibor Hortobágyi, Zsófia Maglóczky, Bernadett Martinecz, Gábor Szabó, Ferenc Erdélyi, Róbert Szipőcs, Benno Gesierich, Marco Duering, István Katona, Arthur Liesz, Gábor Tamás, Ádám Dénes

## Abstract

Microglia are the main immune cells in the brain with emerging roles in brain homeostasis and neurological diseases, while mechanisms underlying microglia-neuron communication remain elusive. Here, we identify a novel site of interaction between neuronal cell bodies and microglial processes in mouse and human brain. Somatic microglia-neuron junctions possess specialized nanoarchitecture optimized for purinergic signaling. Activity of neuronal mitochondria is linked with microglial junction formation, which is rapidly induced in response to neuronal activation and blocked by inhibition of P2Y12-receptors (P2Y12R). Brain injury-induced changes at somatic junctions trigger P2Y12R-dependent microglial neuroprotection, regulating neuronal calcium load and functional connectivity. Collectively, our results suggest that microglial processes at these junctions are in ideal position to monitor and protect neuronal functions in both the healthy and injured brain.

**One-sentence summary:** Neuronal cell bodies possess specialized, pre-formed sites, through which microglia monitor their status and exert neuroprotection.

## Introduction

Microglia are the main immunocompetent cells of the nervous system and their role in brain development and maintenance of proper neuronal function throughout life is widely recognized (*1*–*3*). Importantly, changes in microglial activity are linked with major human diseases including different forms of neurodegeneration, stroke, epilepsy and psychiatric disorders (*4, 5*), directing increased attention towards microglial research in recent years.

Microglia perform dynamic surveillance of their microenvironment by motile microglial processes including their constant interactions with neurons (*6, 7*), yet, the molecular mechanisms of bidirectional microglia-neuron communication have remained elusive. To date, the majority of studies have focused on the interactions between microglial processes and synaptic elements, including axonal boutons and dendritic spines, which have been commonly perceived as the main form of interaction between microglia and neurons (*8, 9*). However, neurons are extremely polarized cells with a high degree of functional independence concerning metabolism and signal integration in their dendritic and axonal compartments (*10*–*12*). In line with this, the large-scale structure of neurons (i.e. their cell body and axonal / dendritic branches) in the brain is relatively stable under most conditions, unlike highly dynamic small synaptic structures, such as dendritic spines and axonal boutons (*13*), which are often distant from neuronal cell bodies. Thus, we argued that the interactions between microglia and synapses may not fully explain how microglia are capable of monitoring and influencing the activity of neurons via dynamic surveillance, or detecting early events of cellular injury in the perisomatic compartment. This may be particularly relevant for the migration and differentiation of neural precursors, cell survival and programmed cell death, adult neurogenesis and the phagocytosis of damaged neuronal cell bodies (*14*–*17*). In line with this, it is not understood how microglia could monitor neuronal status over years or even decades, and discriminate salvageable neurons from irreversibly injured cells mainly based on changes occurring at distant synaptic structures.

To understand the possible mechanisms of effective communication between microglia and neuronal cell bodies, we tested the hypothesis that specialized junctions on neuronal somata may be present that support the dynamic monitoring and assistance of neuronal function by microglia. Here, using *in vivo* two-photon (2P) imaging, high-resolution light- and electron microscopy combined with advanced 3D-analysis we identified a novel type of morpho-functional communication site between microglial processes and neuronal cell bodies in mice and human. Surprisingly, these somatic microglial junctions are present on the vast majority of neurons regardless of their cell type, and have specialized molecular composition and a unique nanoarchitecture linked to mitochondrial signaling. Furthermore, we provide direct *in vivo* evidence that these somatic microglial junctions are essential for microglia-neuron communication, and for the neuroprotective effects of microglia after acute brain injury.

## Results

### Microglial processes contact specialized areas of neuronal cell bodies in the mouse and the human brain

To visualize microglia together with cortical neurons and to study microglia-neuron interactions in the intact brain in real time, CX3CR1^+/GFP^ microglia reporter mice were electroporated *in utero* with *pCAG-IRES-tdTomato* plasmid (Fig. S1a). *In vivo* 2P imaging revealed microglial processes contacting the cell bodies of cortical layer 2-3 neurons in the adult brain (Fig. 1a, b; Movie S1). Microglial processes preferentially returned to the same areas on the neuronal soma, and trajectory-analysis revealed that the average lifetime of somatic microglia-neuron contacts was 25 min, while dendritic contacts had a significantly shorter lifetime of 7.5 min (Fig. 1c). Post-hoc confocal laser scanning microscopy (CLSM) and electron microscopic analysis further validated the direct interaction between microglial processes and the cell bodies of cortical pyramidal neurons (Fig. 1d; Fig. S1c). Similar somatic microglial junctions were present on well-characterized interneuron populations, namely type 3 vesicular glutamate transporter positive (vGluT3+) and parvalbumin-expressing (PV+) cells in the neocortex and the hippocampus (Fig. 1e). Importantly, CLSM revealed the presence of somatic microglia-neuron junctions in the human neocortex as well (Fig. 1f). Quantitative analysis uncovered that somatic microglial junctions were present on 91% of cortical pyramidal cells, 96% of vGluT3+, and 87% of PV+ interneurons in mice. Despite the well-established microglial regulation of neuronal synapses, only 9% of glutamatergic and 14% of GABAergic synapses were associated with microglial processes (Fig. 1g; Fig. S1d). Remarkably, 87% of neurons in the human neocortex were found to receive microglial contact onto their cell body (Fig. 1f-g).

**Figure 1.**
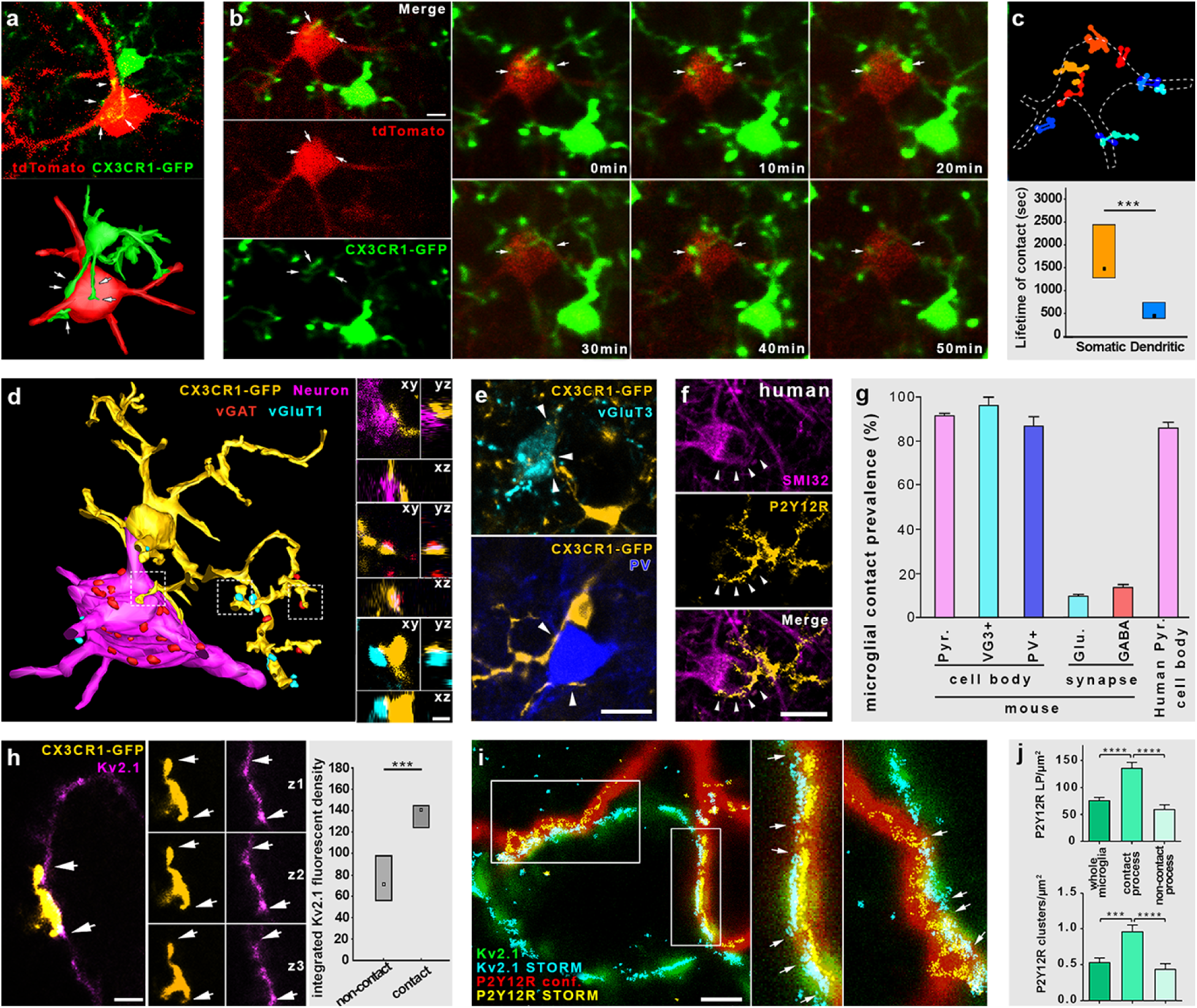
Microglia contact specialized areas of neuronal cell bodies in the mouse and the human brain. a) Single image plane (upper panel) and 3D reconstruction (lower panel) from an in vivo 2-photon (2P) Z-stack shows a neocortical neuron (red) being contacted by microglial processes (green). b) In vivo 2P time-lapse imaging shows temporal dynamics of microglia-neuron contacts. c) Analyzed trajectories of microglial processes contacting the neuron on B. The lifetime of somatic contacts was significantly longer than dendritic ones. (p<0.001, MWU-test, n=26 contacts from 3 mice.) d) 3D-reconstruction from high-resolution confocal laser scanning microscopy (CLSM) Z-stack shows that microglial processes (yellow) contact GABAergic (red) and glutamatergic (cyan) boutons as well as the neuronal cell body (Kv2.1 labeling, magenta). Inserts on the right show orthogonal projections of these contacts from the confocal Z-stack. e) CLSM image planes show yellow microglial processes touching cell bodies of vGluT3+ and PV+ interneurons. f) CLSM images show P2Y12R+ microglial processes contacting SMI32+ neuronal cell bodies in human neocortex. g) Quantitative analysis of contact prevalence between microglial processes and different neuronal elements confirms that microglia contact the vast majority of neuronal cell bodies independently from neurochemical identity, while only a small fraction of synapses receive microglial contact. (n=155 neurons and 800 synapses in mice, and 89 neurons in human). h) CLSM image shows a neuronal cell body contacted by a microglial process. Inserts show three consecutive image planes of the same contact. The integrated fluorescent density of Kv2.1 signal is significantly higher within the contact site than elsewhere. (p<0.001, MWU-test, n=114 ROIs). i) Overlaid images show microglial P2Y12R (red for CLSM and yellow for STORM) and neuronal Kv2.1 (green and cyan) clusters overlapping. Arrows show borders of Kv2.1 clusters. j) P2Y12R clustering depends on the contact with neuronal cell body. Bar graphs show STORM LP density (top) and density of identified P2Y12R clusters (bottom) on different parts of microglia (for statistics, see Table S1). Scale bars: 5 µm on b, 1 µm on d, h, and i, 15 µm on e, 20 µm on f.

Next, we argued that microglia at somatic junctions may sense changes in neuronal state via signals released by exocytosis, which requires a specific molecular machinery supporting membrane trafficking. In neurons, clustered Kv2.1 proteins are well known to form these exocytotic surfaces, anchoring vesicle fusion molecules to the neuronal membrane (*18, 19*). Surprisingly, we found that microglia contacted neuronal somatic membranes at sites of Kv2.1 clustering. The integrated density of Kv2.1 signal at these sites was 95% higher compared to those without microglial contacts (Fig. 1h). The striking association between Kv2.1 clusters and microglial processes was also observed on human cortical neurons (Fig. S1e, f). These Kv2.1 hot spots define preformed neuronal microdomains, because Kv2.1 custering remained unaffected after selective elimination of microglia by PLX5622, confirming their neuron-intrinsic nature (Fig. S1g).

Because activity-dependent exocytotic ATP or ADP release is known to take place from neuronal cell bodies under physiological conditions (*20, 21*) and ATP (ADP) is a major chemoattractant for microglial processes via the microglial purinoceptor, P2Y12R (*6, 22*), we next tested the hypothesis that signaling via P2Y12R is also essential for microglia-neuron interactions at these somatic junctions. In fact, microglia – but no other cells in the brain – were found to be P2Y12R-positive (Fig. S1b), including their processes recruited to somatic junctions (Fig. S1h). To investigate the nanoscale architecture of P2Y12R at somatic microglia-neuron contacts, we used correlated CLSM and STORM superresolution microscopy, which enables the precise assessment of P2Y12R and Kv2.1 clusters at 20 nm lateral resolution (*23*). We found that P2Y12R forms dense clusters on microglial processes at somatic junctions, directly facing neuronal Kv2.1 clusters (Fig. 1i). Importantly, unbiased cluster analysis revealed that P2Y12R localization point density and cluster density were both significantly higher on microglial processes inside the junctions than on processes outside the contacts or on the whole microglial cell (Fig. 1j; Fig. S1i; Table S1). Furthermore, somatic contact-dependent clustering of P2Y12R was found to be a general phenomenon occurring on both pyramidal cells and interneurons (Fig. S1j; Table S1). Contact-dependent molecular clustering, however, could not be observed in the case of the microglial calcium-binding protein, Iba1 (Fig. S1k), suggesting the presence of a functionally specialized, yet ubiquitous communication site between P2Y12R-positive microglial processes and neuronal cell bodies.

### Somatic microglia-neuron junctions possess a uniqe nano-architecture and molecular fingerprints suggesting mitochondrion-related purinergic cell-to-cell communication

To further investigate the ultrastructural features of somatic microglia-neuron junctions, we performed transmission electron microscopy and high-resolution electron tomography with 3D reconstruction. P2Y12R immunogold labeling confirmed the formation of direct junctions between microglial processes and neuronal somata both in mice (Fig. 2a) and in postmortem human brain tissue (Fig. S2a). Surprisingly, we found that microglia-neuron junctions possess a unique ultrastructure within the neuronal cell body composed of closely apposed mitochondria, reticular membrane structures, intracellular tethers and associated vesicle-like membrane structures (Fig. 2a). 3D electron tomography confirmed this nano-architecture in neurons (Fig. 2b; Movie S2-3). These morphological features could not be observed in perisomatic boutons contacted by microglia, confirming the specific association of these structures with somatic microglial junctions. Furthermore, automated 3D analysis of tomographic volumes showed that P2Y12R density negatively correlated with the distance between microglial and neuronal membranes within the junctions (Fig. 2c, d; Fig. S2c), supporting the contact-dependent enrichment of P2Y12Rs on microglial processes. We also compared P2Y12R density between microglial membrane surfaces establishing junctions with neuronal somata and adjacent surfaces (within a few µm-s), that contacted boutons or other neuronal elements. We could detect a significantly higher P2Y12R density at microglial membranes directly contacting neuronal cell bodies (Fig. 2e; Fig. S2d; Movie S4), suggesting an important role for purinergic signaling in the formation of somatic microglia-neuron junctions. Taking advantage of the ultra-high isotropic resolution provided by electron tomography, we could also observe discrete intercellular structures in the extracellular space resembling cell adhesion molecules, connecting the membranes of microglia and neuronal cell bodies (average length 23.5±3.1 nm, Fig. S2b). This falls in the range of the size of integrins expressed by microglia (*24, 25*) or the width of immunological synapses between peripheral immune cells (*26*). Mitochondria-associated membranes (MAM, average distance: 19.5 nm; Fig. S2b) (*27*), and discrete tethers could also be observed between mitochondria and MAM (Movie S3).

**Figure 2.**
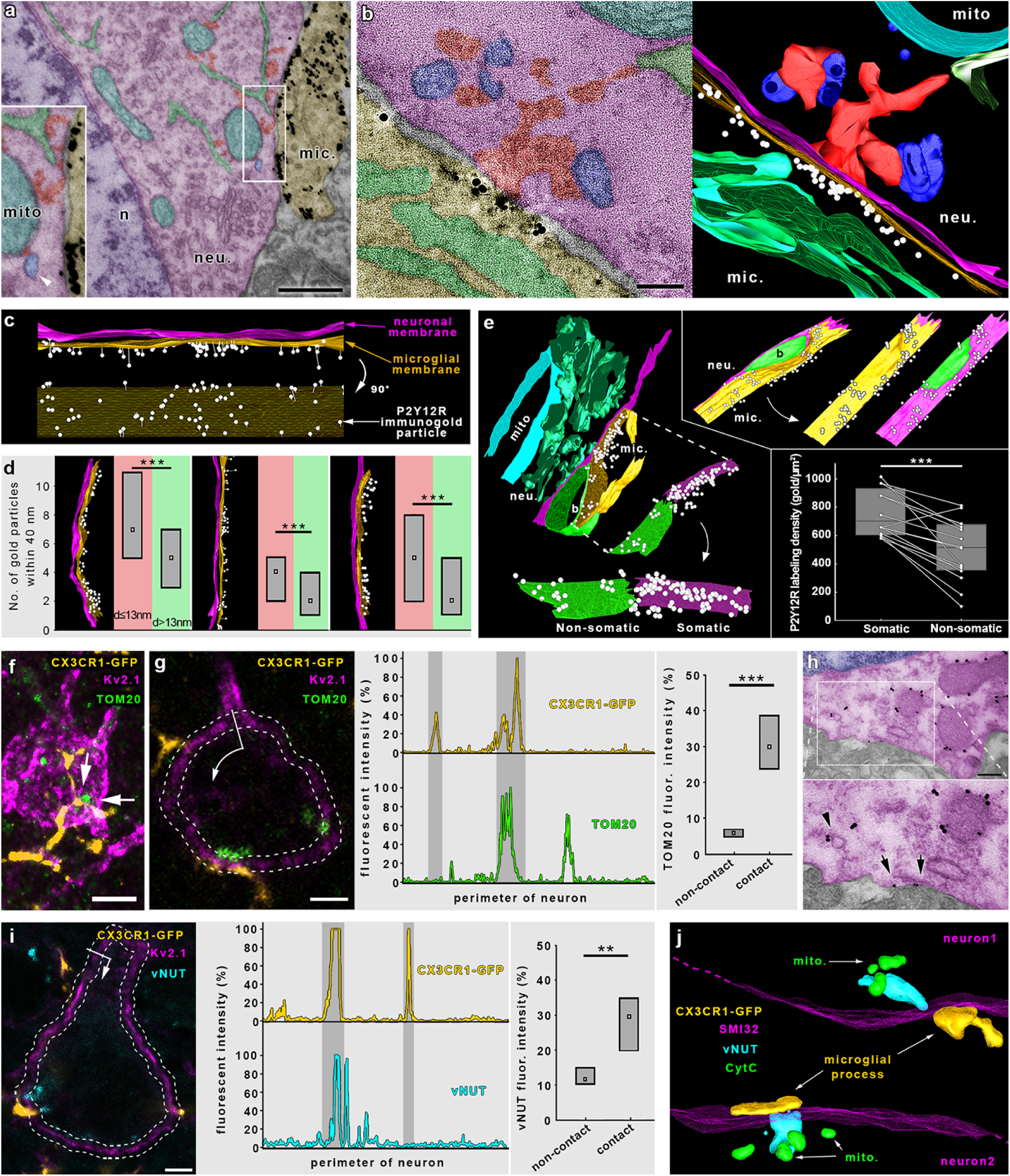
Microglia-neuron junctions possess a specialized nano-architecture and molecular machinery optimized for purinergic cell-to-cell communication. a) Transmission electron micrograph shows the area of the neuronal cell body (neu.) contacted by a P2Y12R-immunogold (black grains) labeled microglial process (mic.). The junction possesses a unique ultrastructure with closely apposed mitochondria (cyan), reticular membrane structures (green), intracellular tethers (red). A mitochondria associated vesicle (blue, marked by white arrowhead) is also visible. b) 0.5 nm thick virtual section of an electron tomographic volume (left) and 3D model (right) shows the special nano-architecture of a somatic microglia-neuron. Note the specific enrichment of P2Y12R labeling at the core of the junction. c-d) P2Y12R density negatively correlates with the distance between microglial and neuronal membranes within the junctions. c) Side-and face (90 degree rotated) view of the 3D model of a contact site. Immunogold density is the highest where the intercellular distance is the smallest. d) Density versus distance analysis performed on 3D models of contact sites confirms that the density of P2Y12R-immunogold particles within the junction is significantly higher where the distance between neuronal and microglial membrane is smaller than 13 nm. (p<0.001, MWU-test, n=13055 points from 3 contacts). e) P2Y12R density is highest at those surfaces of microglial processes that are in direct contact with the neuronal cell bodies. Different views of the 3D models of two junctions illustrate that P2Y12R-immunogold density shows an uneven distribution along microglial membranes, being strongly and selectively enriched where the processes are in direct contact with neuronal somata (mitochondria – mito. neuron – neu., microglia – mic., bouton – b., microglial membrane contacting neuronal cell body – „Somatic”, microglial membrane contacting profiles other than neuronal somata – „Non-somatic”). (p<0.001, MWU-test, n=24 surfaces). f) CLSM maximal intensitiy projection shows microglial processes (yellow) contacting neuronal somata (magenta) with adjacent mitochondria (green). g) Neuronal mitochondria are enriched at microglial junction sites, as TOM20 immunofluorescent intensity is significantly higher along neuronal membrane parts where microglial processes contact the somata. (p<0.001, MWU-test, n=14 contacts). h) Transmission electron micrographs show TOM20-immunogold labeling in neocortical neurons. Immunogold labeling (black grains) is specifically associated with outer mitochondrial membranes, while TOM20-positive vesicles can also be observed (arrowheads). Some immunogold particles can be found on the plasma membrane of the neurons (arrows), suggesting the exocytosis of mitochondria derived vesicles. i) vNUT immunofluorescent intensity is significantly higher in neurons where microglial processes contact the somata. (p=0.002, MWU-test, n=15). j) 3D reconstruction of high-resolution confocal Z-stack shows parts of two neuronal cell bodies (magenta), both contacted by microglial processes (yellow). The vNUT signal (cyan) was concentrated between the junctions and closely positioned mitochondria (green). Scale bars: 500 nm on a, 100 nm on b, 5 µm on f, 2 µm on g, 300 nm on h and 3 µm on i.

Since mitochondrial ATP-production and changes in neuronal activity could trigger microglial process recruitment (*28*–*30*), we investigated the possible enrichment of neuronal mitochondria at microglial junctions on a large sample size, using an unbiased, semi-automatic analysis of TOM20 expression, which is the main element of the transport protein complex in the outer mitochondrial membrane (*31*). TOM20 immunofluorescent intensity was 420% higher at somatic junctions compared to adjacent areas (Fig. 2f, g). Immunoelectron microscopy demonstrated the presence of TOM20-containing vesicles between mitochondria and the neuronal membrane, suggesting the trafficking and possible exocytosis of mitochondria-derived vesicles (*32*) at somatic microglial junctions (Fig. 2h, Fig. S2e). We also found that Kv2.1-immunogold clusters were tightly associated with the observed neuronal structures (i.e. closely apposed mitochondria, MAMs, vesicle-like structures, cytoplasmatic densities) within these junctions (Fig. S2f). Similarly to our CLSM results (Fig. S1g), Kv2.1 nanoclustering was not affected by the absence of microglia (Fig. S2g), confirming the neuron-intrinsic nature of these morpho-functional units, and suggesting that they may function as mitochondria-related signaling hubs in neurons that microglia can specifically recognize. Vesicular release of mitochondria-derived ATP from neurons may occur via vesicular nucleotide transporter (vNUT) (*33, 34*). Indeed, we found that vNUT signal intensity was 2.5 times higher in the vicinity of the neuronal membranes at somatic microglia-neuron junctions than at areas outside the junctions (Fig. 2i). Neuronal vNUT labeling was concentrated between mitochondria and the microglia-contacted neuronal membranes (Fig. 2j). Importantly, Kv2.1 or vNUT signal was not present in perisomatic axon terminals (GABAergic synaptic boutons) including those contacted by microglial processes (Fig. S2h, i), confirming again that these molecular fingerprints are associated specifically with somatic microglia-neuron junctions.

### Physiological microglia-neuron communication at somatic junctions is P2Y12R-dependent and is linked with neuronal mitochondrial activity

Next, we aimed to test whether microglial process recruitment to somatic junctions is functionally linked with the activity of mitochondria in neurons. To this end, CX3CR1^+/GFP^ mice were electroporated with the mitochondria-targeted *CAG-Mito-R-Geco1* reporter construct (Fig. S3a), which further confirmed the involvement of somatic mitochondria in microglial junctions (Fig. 3a). *In vivo* 2P imaging was performed to monitor microglial process recruitment to neuronal mitochondria in the cerebral cortex (Fig. 3b). In line with our histological data, formation of somatic microglial junctions *in vivo* was characterized by the proximity of neuronal mitochondria. Moreover, recruited microglial processes came to close apposition with neuronal mitochondria and stayed there for 15-25 minutes *in vivo* (Fig. 3b, Movie S5). Next, to study the functional relationhsip between microglial junction formation and activity of neuronal mitochondria, we assessed intracellular changes of the metabolic electron carrier nicotinamide adenine dinucleotide (NADH) (*35, 36*) in coronal slices of visual and somatosensory cortices from CX3CR1^+/GFP^ mice. Intracellular NADH fluorescence showed a granular pattern indicating mitochondrial NADH source. Indeed, NADH signal perfectly co-localized with *Mito-R-Geco1* signal, confirming its mitochondrial origin (Fig. 3c). To search for somatic junction formation, we performed 2P imaging, which allowed us to track the movement of microglial processes and monitor cytosolic NADH in viable layer 2/3 neurons simultaneously (Fig. S3b). We could detect apparent increases in NADH intrinsic fluorescence (Fig. 3d, f) parallel with the formation of somatic microglial junctions. In contrast, we found no changes in the mean intrinsic NADH fluorescence detected at neuronal somata contacted by microglial processes in P2Y12R^−/−^ tissue (Fig. 3e, f). These data suggest that microglial process recruitment to somatic junctions is linked to the metabolic activity of neuronal mitochondria via a P2Y12R-dependent mechanism.

**Figure 3.**
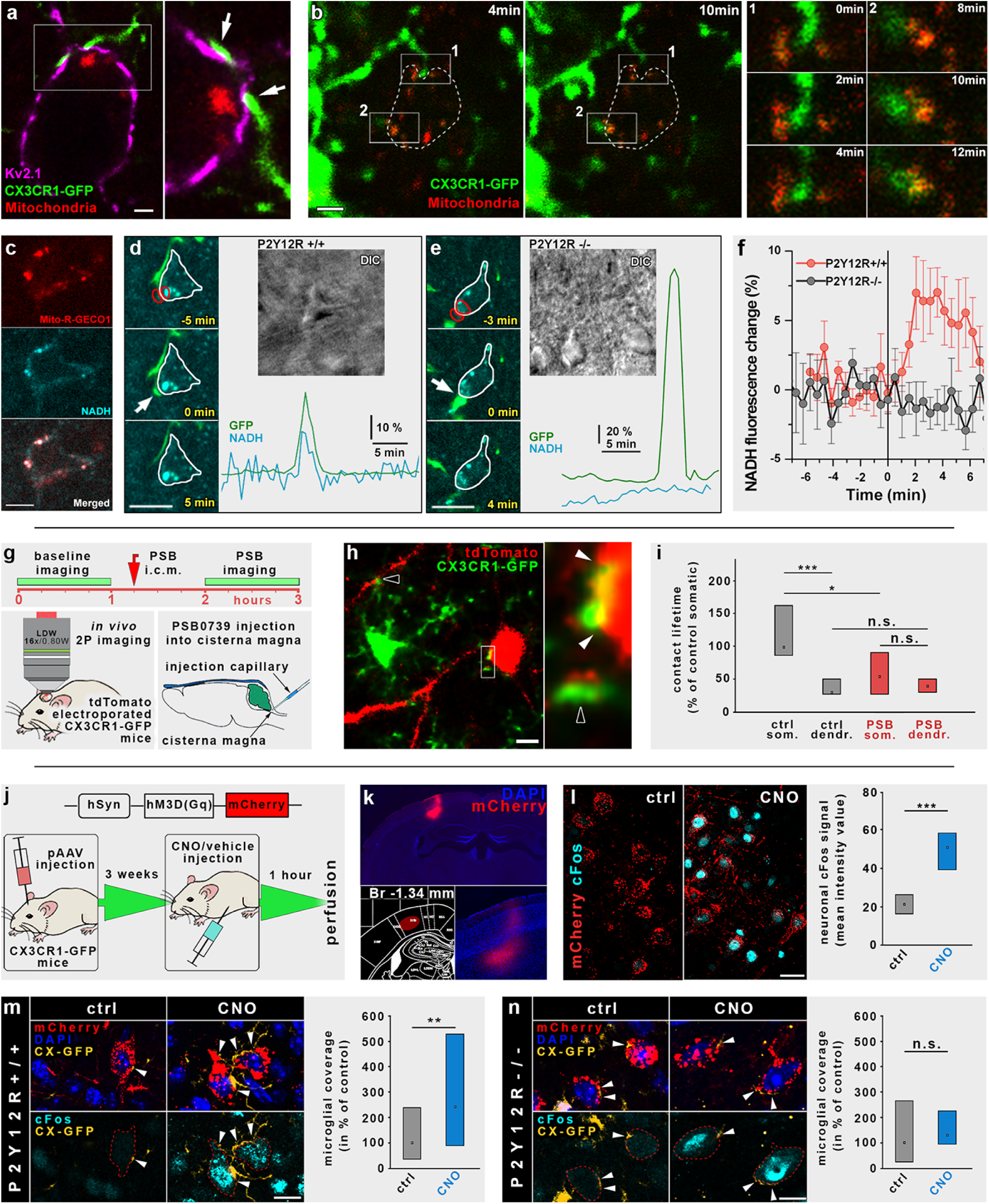
Physiological microglia-neuron communication at the somatic junction site is P2Y12R-dependent and is linked with neuronal mitochondrial activity. a) CLSM image shows a microglial process (green) contacting Kv2.1 clusters (magenta) on a neuronal soma in the vicinity of a mitochondrion (Mito-R-Geco1, red). b) In vivo 2P imaging of CX3CR1^+/GFP^ mice in utero electroporated with CAG-Mito-R-Geco1 construct. Dashed line shows the outline of the neuron, green microglial processes touch neuronal cell body where somatic mitochondria are present. ROIs 1 and 2 are enlarged to show the development of somatic junctions. c) Mito-R-Geco1 expression colocalizes with nicotinamide adenine dinucleotide (NADH) intrinsic fluorescence. d-e) Representative samples from time-lapse imaging of microglia show processes extend and contact neuronal soma in CX3CR1^+/GFP^/P2Y12R^+/+^ (d) and CX3CR1^+/GFP^/P2Y12R^−/−^ (e) mice. White arrow indicates the contact site of microglia. DIC images of the imaged neurons and the fluorescence signal of GFP (green) and NADH (dark cyan) of red outlined areas are shown. f) Average of NADH intrinsic fluorescence of all neurons in P2Y12R^+/+^ (red, n=10) and P2Y12R^−/−^ mice (black, n=11). g) Outline of acute P2Y12R-blockade experiments. Baseline in vivo 2P imaging of cortical microglia-neuron contacts of tdTomato electroporated CX3CR1^+/GFP^ mice was followed by administration of the P2Y12R-inhibitor PSB0739 (PSB) into the cisterna magna (i.c.m.) and a further imaging session. h) Examples of the recorded microglia-neuron contacts. Empty arrowheads point to dendritic contacts, full arrowheads mark somatic junctions. i) Acute i.c.m. administration of PSB significantly reduced somatic junction lifetime, but did not affect the lifetime of dendritic microglia-neuron contacts (ctrl som. vs. PSB som. p=0.0331, MWU-test, n=40). j) Outline of chemogenetic experiments. k) An example of an injection site. l) CNO administration induced a 2.3-fold increase in cFos mean fluorescent intensity value in DREADD expressing neurons (p<0.001, MWU-test, n=100). m) Neuronal activity induced a robust elevation of microglial process coverage of neuronal cell bodies in CNO-treated animals (p=0.0055, MWU-test, n=85 cells). n) CNO triggered neuronal activity could not induce an elevation of microglial process coverage of neuronal cell bodies in P2Y12R^−/−^ mice (p=0.26, MWU-test, n=65 cells). Scale bars: 2 µm on a, 4 µm on b, 5 µm on c and h, 10 µm on d and e, 8 µm on m and n.

Because our data strongly suggested a role for somatic junctions in sensing neuronal activity by microglia, we aimed to further explore the signaling mechanisms at these sites *in vivo* by 2P imaging in CX3CR1^+/GFP^ microglia reporter mice that were electroporated *in utero* with the neuronal reporter *pCAG-IRES-tdTomato* (Fig. 3g-h). Administration of the potent and selective P2Y12R-inhibitor PSB0739 into the cisterna magna (i.c.m.) significantly reduced somatic junction lifetime by 45%, but did not affect the lifetime of dendritic microglia-neuron contacts (Fig. 3g, i). Since these data showed that the maintenance of somatic microglia-neuron junctions depends on physiological P2Y12R function, we aimed to test whether microglia would react directly to changes in neuronal activity. To this end, we induced neuronal activation by using the chemogenetic DREADD (Designer Receptor Exclusively Activated by Designer Drug) approach. pAAV carrying the *hSyn-hM3D(Gq)-mCherry* construct was injected into the cerebral cortex of P2Y12R^+/+^ and P2Y12R^−/−^ mice that had been crossed with CX3CR1^+/GFP^ mice to visualize microglial responses in the presence or absence of P2Y12R signaling (Fig. 3j, k). After intraperitoneal injection of clozapine N-oxide (CNO) to induce hM3D(Gq)-DREADD activation, we observed a 250% increase in neuronal cFos signal compared to vehicle treatment (Fig. 3l), confirming a specific and robust neuronal activation. We found that chemogenetic neuronal activation resulted in an increased microglial process coverage of the soma of DREADD-expressing neurons in P2Y12R^+/+^ mice (Fig. 3m), but not in P2Y12R^−/−^ mice (Fig. 3n).

Collectively, these results confirmed that microglia dynamically react to changes in neuronal activity at somatic microglia-neuron junctions in a P2Y12R-dependent manner, leading to a rapid increases of somatic coverage by microglial processes.

### Microglia protect neurons after acute brain injury in a P2Y12R-dependent manner via altered somatic junctions

Since somatic microglia-neuron junctions were abundant in the healthy brain, we next examined how these morpho-functional communication sites are altered in response to brain injury. Microglia are known to rapidly respond to changes in neuronal activity in the boundary zone of the infarct after stroke (*37*). Thus, we performed experimental stroke and delineated the evolving penumbra based on the metabolic activity of the tissue as assessed by the redox indicator TTC (tetrazolium chloride) coregistered with the immunofluorescent signal for MAP2 and microglia (Fig. S4a). We observed the fragmentation of mitochondria and an almost complete declustering of Kv2.1 proteins (*38, 39*) in morphologically intact penumbral neurons (Fig. 4a, b; Fig. S4e). These morphological changes were accompanied by a robust (380%) increase in the microglial process coverage of neuronal cell bodies, originating from somatic microglia-neuron junctions in both mice and human post-mortem brain tissues (Fig. 4b-e). Strikingly, acute i.c.m. administration of the P2Y12R-inhibitor PSB0739, or preventing mitochondrial injury by using the mitochondrial ATP-sensitive potassium (KATP) channel opener diazoxide (*40, 41*), completely abolished stroke-induced increases in microglial process coverage around somatic junctions (Fig. 4d). Viability of the examined neurons with increased microglial process coverage was confirmed by normal chromatin structure and membrane integrity (Fig. S4b, c). Transmission electron tomography also confirmed increased microglial process coverage and mitochondrial fragmentation of neurons (Fig. 4c).

**Figure 4.**
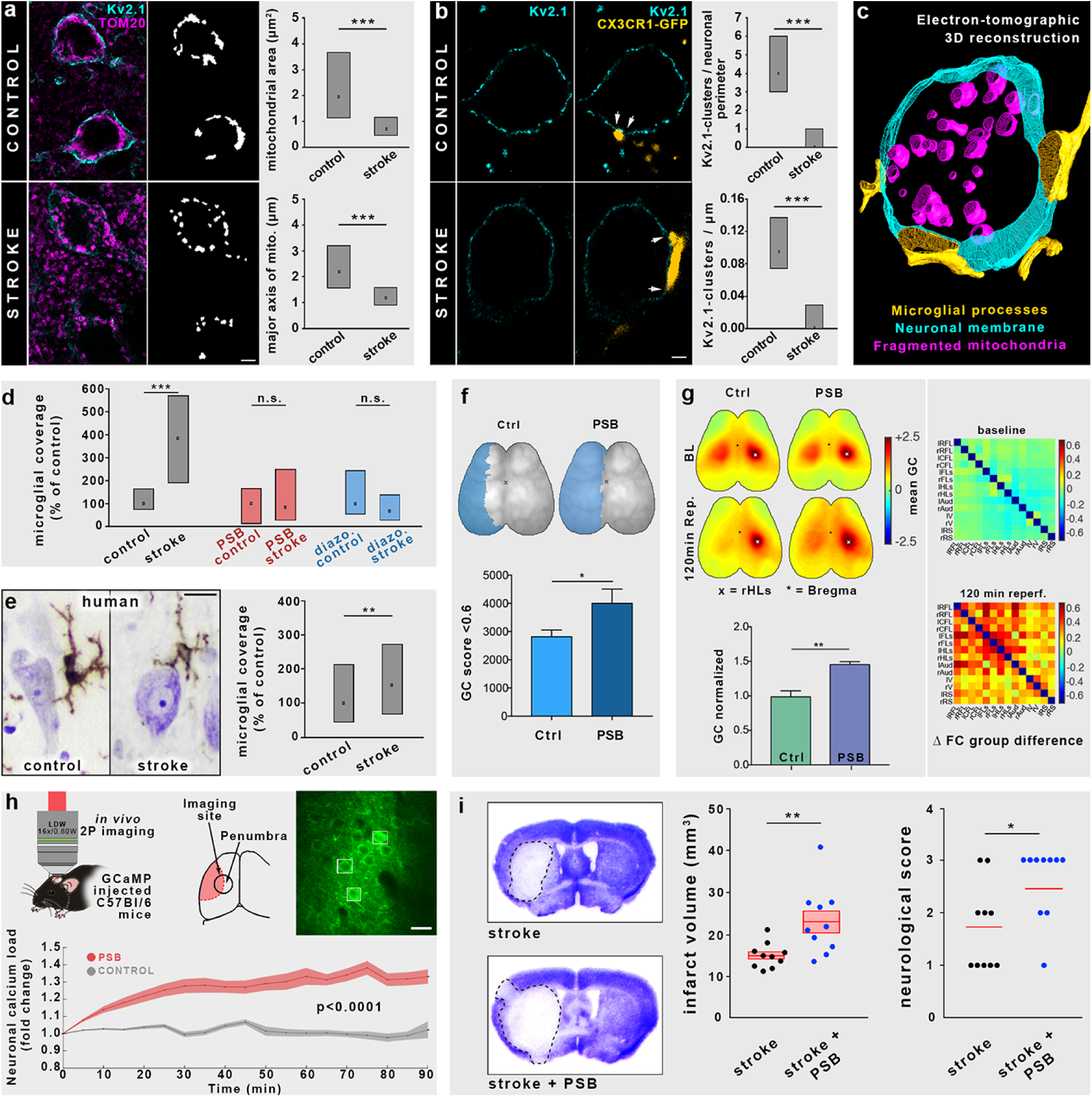
Microglia protect neurons after acute brain injury in a P2Y12R-dependent manner via altered somatic junctions. a) CLSM images show that stroke induces the fragmentation of mitochondria (magenta) in neuronal cell bodies (Kv2.1 labeling, cyan) in the penumbra. Mitochondrial area and mitochondrial major axis are both significantly decreased (p<0.001, MWU-test, n=189 mitochondria). b) CLSM images of cortical neurons show that in parallel with the declustering of Kv2.1-channels (cyan) microglial coverage (yellow) is significantly increased after stroke in the penumbra (p<0.001, MWU-test, n=30 neurons). c) 3D-reconstruction from electron tomographic volume shows elevated microglial coverage and fragmentation of neuronal mitochondria. d) Microglial coverage of neuronal cell bodies is robustly increased after stroke, while acute central blockade of P2Y12Rs or activation of mitochondrial ATP-sensitive potassium (KATP) channels completely abolishes the stroke-induced increase of coverage (ctrl vs. stroke: p<0.001, PSB ctrl vs. PSB stroke: p=0.792, diazo. ctrl vs. diazo. stroke: p=0.053, MWU-test, n=140 neurons). e) Stroke induces a 1.5-fold increase in somatic microglia coverage of human cortical neurons (p=0.007, MWU-test, n=249 neurons). f) Topographical maps show the area of pixels with a global connectivity (GC) score less than 0.6 after during ischemia. The sum of outlined pixels revealed higher dropdown of GC in PSB treated animals after stroke (p=0.0439, unpaired-t-test, n=17 mice). g) Left panel: Topographical maps show increased ROI-to-global connectivity of the contralateral HLs in PSB-treated mice 120 min after stroke. Individual ROI-to-global connectivity scores were normalized to baseline and quantified for group comparison (p=0.0077, unpaired-t-test, n=7 mice). Right panel: Seed-to-seed connectivity is increased in PSB-treated animals after stroke. h) In vivo 2P calcium imaging reveals a significant increase of neuronal calcium load during reperfusion after acute P2Y12R inhibition with PSB (p<0.0001, Kruskal-Wallis test, n=96 neurons). i) Infarct volume is increased after acute central P2Y12R-inhibition, which is accompanied by a significantly worse neurological outcome. Scale bars: 4 µm on a, 2 µm on b, 8 µm on e, 20 µm on h.

To test the impact of P2Y12R-dependent microglial functions on neuronal viability *in vivo*, we investigated pharmacological inhibition of P2Y12R by injection of PSB0739 i.c.m. prior to MCAo. Inhibition of microglial P2Y12R not only prevented increases in microglial process coverage of neuronal cell bodies in the penumbra, but it also altered functional connectivity in the brain as assessed by a widefield imaging approach in Thy1-GCaMP6s mice (Fig. 4f-g). In fact, an absence of P2Y12R signaling significantly increased the area of functional disconnection (global connectivity < 0.6) in the ipsilateral hemisphere during ischemia, accompanied by a trend towards elevated neuronal calcium-load (Fig. 4f; Fig. S4f). Seed-based connectivity analysis revealed a significant increase in the contralateral sensory hindlimb area after reperfusion in PSB0739 treated animals. Moreover, connectivity analysis of 14 functional areas revealed a substantial and widespread increase in connectivity strength in the absence of microglial P2Y12R signaling (Fig. 4g). To examine the effect of P2Y12R inhibition at the single neuron level in the evolving ischemic penumbra *in vivo*, we investigated GCaMP6f-injected mice with 2P microscopy. In control mice, neuronal GCaMP6f signal remained unchanged for the first 90 minutes of reperfusion, while blockade of microglial P2Y12Rs with PSB0739 resulted in a strong elevation in neuronal calcium load (Fig. 4h), corroborating the findings obtained from the widefield imaging approach also at the cellular level. Most importantly, P2Y12R inhibition significantly increased lesion volume at 24 h reperfusion (Fig. 4i) and led to worse neurological outcome (Bederson score, Fig. 4i). Thus, the above data collectively show that disintegration of somatic microglia-neuron junctions after neuronal injury triggers increased microglial process coverage of the cell bodies of compromised but potentially viable neurons via P2Y12R and mitochondrial signaling, allowing the initiation of protective microglial responses that limit brain injury.

## Discussion

Here we identify a novel form of interaction between microglia and neurons. We show that under physiological conditions, somatic microglia-neuron junctions are present on the majority of neurons in both mice and humans, function as major communication sites and are rapidly altered in response to brain injury. We propose that microglia constantly monitor neuronal status via these newly identified somatic junctions, allowing neuroprotective actions to take place in a targeted manner.

Our *in vivo* 2P imaging, CLSM and electron microscopic studies showed that sites of somatic junctions in neurons are preferentially and repeatedly contacted by microglia and such interactions have profoundly increased lifetime compared to the microglial contacts targeting dendrites. In previous studies, the proximity between microglial cell bodies or processes with neuronal somata have been observed in zebrafish and mice (*42, 43*). However, activity-dependent recruitment of microglial processes to neuronal cell bodies, the formation of direct membrane-to-membrane junctions, the molecular identity of neuronal membranes contacted, the mechanisms of junction formation, and the function of somatic microglia-neuron interactions have not been experimentally addressed. Thus, we took advantage of cutting edge neuroanatomical approaches that have not been previously used to study microglia-neuron interactions, and discovered that somatic microglia-neuron junctions are characterized by unique ultrastructural and molecular composition. These morphological and molecular features are absent in perisomatic boutons contacted by microglia (Fig. S2h-i), suggesting that the main form of neuronal quality control by microglial processes is not mediated by interactions between microglia and perisomatic axon terminals.

Mitochondria are the primary energy generators in cells, playing fundamental roles in calcium homeostasis, intracellular signaling (*28, 30, 44*), neuronal quality control (*45*), and in determining cellular fate (*46, 47*). While neuronal mitochondria are also considered as “immunometabolic hubs” involved in antigen presentation and the regulation of innate immune responses (*48, 49*), changes in mitochondrial function due to metabolic imbalance, oxidative stress, inflammation, cellular injury or cell death occur in most neuropathological states (*50*). Mitochondria-associated membranes (MAM) are also considered to be key integrators of metabolic and immunological signals, playing a central role in neurodegeneration and cell-fate decisions (*27, 51*–*53*). Thus, somatic mitochondria and MAMs are ideally positioned to report neuronal status to microglia and to mediate neuronal quality control. In line with this, we show that the recruitment of microglial processes to somatic junctions in the vicinity of neuronal mitochondria is linked with mitochondrial activity (NADH fluorescence) changes in neurons via P2Y12R signaling. Neurons can execute somatic ATP release via pannexin hemichannels, voltage dependent anion channels or through activity-dependent vesicle exocytosis (*20, 21, 54*). Vesicular nucleotide transporter (vNUT) is known to be responsible for somatic vesicular ATP-release in neurons (*34*). In fact, we demonstrated the enrichment of vNUT between neuronal mitochondria and the somatic membranes contacted by microglia, where TOM20-positive mitochondria-derived vesicles were also observed, through which regulated exocytotic release of ATP and other vesicular substances could provide a constant readout of neuronal activity and mitochondrial function as seen in neurons and other cells (*32, 55, 56*). The strong enrichment of vNUT in these contacts, the presence of filamentous cytoplasmatic structures connecting vesicles to the core of the junction (Fig. 2a-b), the presence of TOM20 immunogold particles on the neuronal plasmamembrane (Fig. 2h) and the massive accumulation and nanoscale clustering of exocytosis-promoting Kv2.1-proteins within these contact sites collectively indicate exocytotic vesicular release at microglia-neuron junctions, which will need to be investigated in future studies.

Kv2.1 channels are major regulators of neuronal potassium levels, however, they tend to assemble into discrete clusters on the surface of neurons, where they do not function as ion channels, but provide sites for intensive membrane trafficking as exo- and endocytotic hubs (*18, 57, 58*). Furthermore, Kv2.1 clusters are known to induce stable ER-plasmamembrane junctions (*58*), localizing MAMs and mitochondria into these morpho-functional units (Fig. S2f), providing an ideal site for release of mitochondria-associated messenger molecules (*32*). Neuronal activity or noxious stimuli lead to the rapid dispersion of Kv2.1-clusters (*39, 59, 60*). We found that Kv2.1 declustering takes place in compromised neurons of the penumbra as early as 4 hours after brain injury that paralleled mitochondrial fragmentation in neurons, and increased microglial process coverage around somatic microglia-neuron junctions. Our present studies did not investigate whether Kv2.1 declustering directly shapes microglial responses, or Kv2.1 signal simply outlines changes in somatic junction structure, indicative of mitochondrial dysfunction that triggers an increase of microglial process coverage of neurons and the initiation of protective microglial responses via additional molecular mechanisms. However, using STORM super-resolution imaging, we found that microglial P2Y12R-clusters are precisely aligned with neuronal Kv2.1 clusters at somatic junctions (Fig. 1i). Interestingly, the activation of P2Y12R-s were mainly associated with injury or pathological states in previous studies, and were considered negligable for physiological microglial surveillance, based on *ex vivo* studies (*61*). Our *in vivo* results challenge this view, since the lifetime of somatic microglia-neuron junctions (but not dendritic contacts) and neuronal mitochondrial activity were both dependent on physiological microglial P2Y12R activity. The contact-dependent clustering of these receptors further confirms their involvement in physiological microglia-neuron interactions. Interestingly, blockade of microglial P2Y12R left cortical synapse numbers completely unchanged (Fig. S4g) and contact-dependent nano-clustering of microglial P2Y12R-s at somatic junctions was not seen when microglia contacted synaptic boutons. These structural and functional data collectively suggest that microglia-neuron interactions at these newly identified somatic junctions are not only P2Y12R-dependent, but are fundamentally different from those seen at synapses.

The failure of most neuroprotection trials in stroke and other brain diseases strongly indicates the importance of understanding the complexity of pathophysiological processes, including microglial actions. In fact, potentially salvageable neurons around the infarct core may show metabolic activity up to 6-17 hours following stroke in patients and experimental animals (*62*–*64*). Concerning the fate of these neurons, both the detrimental role of inflammatory mechanisms and the protective nature of microglial actions have been previously demonstrated (*37, 65, 66*). Our present results reveal that P2Y12R-dependent microglial actions protect neurons, while blockade of microglial P2Y12R signaling alone impairs cortical network function, increases calcium load, and the area of ischemia-induced disconnection within two hours following stroke (a clinically relevant time window), leading to augmented brain injury, similarly to that seen after the complete and selective elimination of microglia (*37*). We also show that these protective microglia- and P2Y12R-mediated effects are linked with mitochondrial actions initiated upon neuronal injury, since the KATP-channel opener diazoxide abolished the increases in microglial process coverage of neurons after stroke, similarly to the blockade of P2Y12R signaling.

## Clinical significance

Genome-wide association studies, clinical data and experimental research collectively identify both microglia and neuronal mitochondria as key effectors in common neurological diseases including dementia, psychiatric disorders, acute brain injury and different forms of neurodegeneration (*4, 5, 50, 68*). The exact mechanisms underlying these conditions are currently unclear and effective therapies are lacking, despite all research efforts and numerous clinical trials performed to protect injured neurons in the brain. Based on the available data, microglia-mediated effects in health and disease cannot be explained merely by microglia-synapse interactions that often occur at a great distance from injured neuronal somata, where the majority of mitochondria critical for cell fate decisions are localized (*46*). To resolve these controversies, we describe a novel form of intercellular interaction through which microglia recognize neuron-intrinsic somatic signaling hubs, dynamically monitor neuronal state and mitochondrial function, respond to changes in neuronal activity through increased coverage of neuronal cell bodies and influence neuronal fate in a P2Y12R-dependent manner. These unique somatic microglia-neuron junctions exist in both mice and human in the majority of neurons and are altered in response to brain injury, suggesting their importance. These structures are likely to accommodate presently unexplored molecular fingerprints and signaling pathways involved in intercellular communication, which must be investigated in upcoming studies. We propose that healthy neurons may constitutively release signaling molecules at these junctions, reflecting their “well-being” towards microglia. In turn, disintegration of these specialized morpho-functional hubs due to excitotoxicity, energy depletion or other noxious stimuli may trigger rapid and inherently protective microglial responses, leading to the restoration of neuronal function or isolation and phagocytosis of dying neurons in case terminal neuronal injury occurs (*67*).

The most important open questions include the clarification of additional signaling mechanisms (vesicular and non-vesicular) involved in neuron-to-microglia communication at these junctions, and the mechanisms of microglial neuroprotection (e.g. regulation of neuronal ion-fluxes, or the metabolism of neuronal mitochondria). Since the role of microglia-neuron somatic junctions in most brain diseases is completely unknown, we assume that microglia-neuron interactions through these sites markedly differ in different forms of acute and chronic neuropathologies and have enormous therapeutic potential.

## Author contributions

C.C., B.P. and A.D. conceived the project. Surgery was performed by N.L. and A.D.; two-photon imaging was performed by R.F., immunohistochemistry and light microscopy was performed by C.C., B.P., A.D., B.O., and A.D.S; STORM microscopy was performed by B.O., Electron microscopy was performed by C.C., B.P. and A.D.S., electron tomography was performed by C.C. and B.P., *in vitro* NADH imaging was performed by G.M. under the supervision of G.T., plasmid engineering and *in utero* electroporation was performed by Z. L. and Z. I. L., virus injection was performed by R. F. and B.M., widefield calcium imaging was performed by S.H. under the supervision of A.L.; C.C., B.P., B.O., G.M., S.H., N.L., A.D.S., K.U., A.L., A.D. analysed data, L.C., T.H., Z.M. G.S. and F.E. contributed with critically important materials, R.S. optimized the laser-optical setup for dual wavelength *in vitro* 2P-merasurements, I.K., G.T. and A.L. provided resources and essential intellectual contribution, and revised the manuscript. A.D. obtained funding and supervised the project. C.C. B.P. and A.D. wrote the paper with input from all authors.

## Acknowledgements

This work was supported by „Momentum” research grant from the Hungarian Academy of Sciences (LP2016-4/2016 to A.D.) and ERC-CoG 724994 (A.D.). Additionally, this work was funded by Hungarian Academy of Sciences (G.T.), the National Research, Development and Innovation Office of Hungary (GINOP-2.3.2-15-2016-00018, VKSZ-14-1-2015-0155, G.T.), the Ministry of Human Capacities, Hungary (grant 20391-3/2018/FEKUSTRAT, G.T.), by the Munich Cluster for Systems Neurology (EXC 2145 SyNergy) and ERC-StG 802305 to A.L.; I.K. was supported by „Momentum” research grant from the Hungarian Academy of Sciences (LP2013-54), Hungarian Scientific Research Fund (OTKA, K 116915) and National Research, Development and Innovation Fund (VKSZ_14-1-2015-0155); Zs.M. was supported by National Research, Development and Innovation Office of Hungary (grant K 125436) and by National Brain Research Program (2017-1.2.1-NKP-2017-00002). We thank László Barna and the Nikon Imaging Center at the Institute of Experimental Medicine for kindly providing microscopy support, David Mastronarde at MCDB for his continous help with IMOD software, and Solt Kovács from ETH Zurich for scripting analytic tools. We thank the Department of Pathology, St. Borbála Hospital, Tatabánya, and the Human Brain Research Lab at the Institute of Experimental Medicine, Hungarian Academy of Sciences (IEM HAS) for providing human brain tissue and Dóra Gali-Györkei for her excellent technical assistance. We also thank for Plexxikon for providing PLX5622.

## Methods

### Ethical considerations

All experiments were performed in accordance with the Institutional Ethical Codex and the Hungarian Act of Animal Care and Experimentation guidelines (40/2013, II.14), which are in concert with the European Communities Council Directive of September 22, 2010 (2010/63/EU). The Animal Care and Experimentation Committee of the Institute of Experimental Medicine of Hungarian Academy of Sciences and the Animal Health and Food Control Station, Budapest, have also approved the experiments under the number PE/EA/2553-6/2016. Control human brain tissue was obtained from two female (59- and 60-years-old) and one male (73-years-old) subjects who died from causes not linked to brain diseases, and did not have a history of neurological disorders (ETT TUKEB 31443/2011/EKU [518/PI/11]). Tissues from patients who died after ischemic stroke affecting the MCA area were obtained from two female (77- and 78-years-old) and one male (66-years-old) subjects (ethical approval ETT-TUKEB 62031/2015/EKU, 34/2016 and 31443/2011/EKU (518/PI/11)) See Supplementary Table 2. Informed consent was obtained for the use of brain tissue and for access to medical records for research purposes. Tissue was obtained and used in a manner compliant with the Declaration of Helsinki.

### Animals

Experiments were carried out on 12-18 weeks old C57BL/6 (RRID:IMSR_JAX:000664), CAMK2 GFP, PVA GFP, GAD65 GFP, CX3CR1^+/GFP^ (IMSR_JAX:005582), CX3CR1^+/GFP^/P2Y12^−/−^ and C57BL/6J-Tg(Thy1-GCaMP6s)GP4.12Dkim/J mice (*22, 69*–*72*). Mice were bred and genotyped at the SPF unit of the Animal Care Unit of the Institute of Experimental Medicine (IEM HAS, Budapest, Hungary) as described earlier (*37*). Mice had free access to food and water and were housed under light-, humidity- and temperature-controlled conditions. All experimental procedures were in accordance the guidelines set by the European Communities Council Directive (86/609 EEC) and the Hungarian Act of Animal Care and Experimentation (1998; XXVIII, section 243/1998), approved by the Animal Care and Use Committee of the IEM HAS. All experiments were performed in accordance with ARRIVE guidelines.

### Perfusion, tissue processing for histology

Adult mice were anesthetized by intraperitoneal injection of 0.05-0.1 ml of an anaesthetic mixture (containing 8.3mg/ml ketamine, 1.7 mg/ml xylazin-hydrochloride). Animals were perfused transcardially with 0.9% NaCl solution for 1 minute, followed by 4% freshly depolymerized paraformaldehyde (PFA) in 0.1 M phosphate buffer (PB) pH 7.4 for 40 minutes, and finally with 0.1 M PB for 10 minutes to wash the fixative out. Blocks containing the primary somatosensory cortex and dorsal hippocampi were dissected and coronal sections were prepared on a vibratome (VT1200S, Leica, Germany) at 20 µm thickness for STORM experiments, 50 µm thickness for immunofluorescent experiments and for electron microscopy/electron tomography.

### Post-mortem human brain tissues

Brains of patients who died in non-neurological diseases were removed 4-5 h after death. The internal carotid and the vertebral arteries were cannulated, and the brain was perfused first with physiological saline (using a volume of 1.5 l in 30 min) containing heparin (5 ml), followed by a fixative solution containing 4% paraformaldehyde, 0.05% glutaraldehyde and 0.2% picric acid (vol/vol) in 0.1 M PB, pH 7.4 (4–5 l in 1.5–2 h). The cortical and hippocampal samples were removed from the brain after perfusion, and was postfixed overnight in the same fixative solution, except for glutaraldehyde, which was excluded. Blocks were dissected, and 50 µm thick sections were prepared on a vibratome (VT1200S, Leica, Germany). Brains of stroke patients were removed after 10-15 h after death and immersion fixed in 4% paraformaldehyde. Small regions from the affected cerebral cortex were dissected, embedded into paraffin both from the ipsilateral and contralateral hemisphere and 6-8 µm thick sections were cut on a sledge microtome.

### Cloning

*CAG-IRES-tDTomato* (pCAG-tDTomato): The GFP-polyA part of the pCAGIG plasmid (a gift from Connie Cepko, Addgene plasmid # 11159, RRID:Addgene_11159 (*73*)) was replaced with tDTomato-pA (a gift from Gyula Balla, IEM-HAS, Hungary) using blunt end cloning (pCAGIG:PstI-BstXI, pcDNA3-tDTomato: HindIII-PvuII), they were blunted via Klenow chewback/fill-in respectively.

*CAG-Mito-R-GECO: CMV-Mito-R-GECO1* (a gift from Robert Campbell; Addgene plasmid # 46021, RRID:Addgene_46021 (*74*)) was digested PmeI and the mito-R-Geco fragment was subcloned into EcoRV-digested pBSKII SK+ (Stratagene). Orientation was checked with restriction analysis. Then pBSKII-Mito-R-GECO was cut with Acc65I-NotI and cloned into pCAG-GFP (a gift from Connie Cepko (Addgene plasmid # 11150, RRID:Addgene_11150 (*73*)) digested with Acc65I-NotI.

### Cell culture transfection

HEK-293 cells were cultured in Dulbecco’s Modified Eagle Medium (4.5 g/L glucose, L-glutamine & sodium pyruvate; Corning) with 10% heat inactivated Fetal Bovine Serum (Biosera) and incubated at 37 °C in 5% CO2 in air. On the day of transfection, cells were plated on poly-D-lysine (Sigma) coated 18 mm coverslips in 12-well cell culture plates. For transfection 2 μl Lipofectamine® 2000 Reagent (Thermo Fisher Sc.) were mixed with 2 μg Mito-R-Geco1 in Gibco® Opti-MEM™ Media (Thermo Fisher Sc.) and stored in the hood approximately half an hour. Transfection solution was mixed with the culturing media and cells were incubated overnight. Next day, cells were fixed with 4% PFA for 10 minutes, than washed with PBS. Permeabilization and blocking steps were performed by 0,1% TritonX and 5% NDS (Normal Donkey Serum; Sigma) /PBS solution for 30 mins. TOM-20 antibody (1:1000) was applied in PBS for 90 mins. After several PBS washes cell were treated with secondary antibodies/PBS solution for an hour. Finally coverslips were washed in PBS and mounted with VECTASHIELD® HardSet™ mounting medium and sealed with nail polish. Confocal images were taken with 60X objective by Nikon A1R confocal system guide by NIS-Elements Microscope Imaging Software.

### *In utero* electroporation

Timed-pregnant C57Bl/6J (Jackson) females bred with homozygous CX3CR1^GFP/GFP^ transgenic animals were anesthetized by isoflurane vaporization at embryonic day 14.5. Abdominal cavity was opened longitudinally and uterine horns were exposed. Approximately 1μl of expression vector (1 μg/μl all of the constructs) in endotoxin-free water containing Fast Green dye (Roth 1:10000 dilution) was injected into the embryonic lateral ventricles, using glass capillary and mouth pipette. Electroporation was performed with tweezer electrodes, 5×50 V pulses of 50 millisecond duration was applied with 950 millisecond intervals using the In Utero Electroporator SP-3c (Supertech). After the electroporation, uterine horns were returned into the abdominal cavity, the muscle wall and skin were sutured, and embryos were allowed to continue their normal development. All the littermates were CX3CR1^+/GFP^ heterozygous and were born naturally.

### Pharmacological treatments and chemogenetics

30 min before middle cerebral artery occlusion (MCAo) surgeries a single dose (0,6 mg/kg dissolved in phosphate buffer or saline) of P2Y12 receptor (P2Y12R) antagonist (PSB0739 Tocris, R&D Systems, Minneapolis, USA) was administered to the cisterna magna in 5 µl final volume using a glass capillary. Brains were harvested 24 h later for histology. Diazoxide (#D9035, Sigma, Merck KGaA, Darmstadt, Germany) dissolved in 0,4% DMSO and 0,01 M NaOH was administered in a single dose of 10 mg/kg intra peritoneally, immediately before reperfusion of the MCAo took place during the stroke surgeries. Brains were prepared 4 h later for histological assessment. For selective microglia elimination C57BL/6J mice were fed a chow diet containing the CSF1 receptor antagonist, PLX5622 (Plexxikon Inc., 1200 mg PLX5622 in 1 kg chow) for 3 weeks to eliminate microglia from the brain (and another group of C57BL/6J mice were fed control chow diet). For the chemogenetic activation of neurons 0.1 µl of AVV8-pAAV-hSyn-HA-hM3D(Gq)-MCherry (RRID:Addgene_50474, Addgene, USA) was injected into the neocortex of CX3CR1^+/GFP^ mice. After 3 weeks incubation, mice received saline or clozapine-N-oxide (0.1 mg/ml) intraperitoneally to induce DREADD activation. Animals were perfused 1 hour after the injections, and processed for histology. For *in vivo* calcium imaging C57Bl/6 mice were used. 200 nl concentrated AAV1.Syn.GCaMP6f.WPRE.SV40 (RRID:Addgene_100837, Penn Vector Core) was injected into the cortex 200-300 µm below the surface with glass capillary. The injection coordinates were 1.5 mm lateral from midline, 1.2 mm posterior from bregma. Cranial window surgery and two-photon (2P) resonant imaging were performed 2 weeks after injection.

### Experimental stroke

MCAo was performed using the intraluminal filament technique as described earlier (*75*). In brief, animals were anaesthetized with isoflurane and a silicone-coated monofilament (210-230 µm tip diameter, Doccol, Sharon, US) was introduced into the left external carotid artery and advanced along the internal carotid artery to occlude the MCA for 30 or 60 min. Occlusion was confirmed by a laser Doppler (Moor Instruments, UK). During surgery, core temperature was maintained at 37±0.5°C. The following exclusion criteria were set up: animals having less than 70% relative reduction in blood flow, either having haemorrhage or having shorter survival than 24 h were excluded from any further analysis pre hoc. In total 2 animals (one control and one PSB treated) were excluded post hoc due to intracerebral haemorrhage, 1 PSB treated animal died before 24 h. Altogether 3 out of 36 animals were excluded (28.26%) from the analysis, and total mortality was 2.77%.

Functional outcome of mice was assessed 24 h after MCAo using the corner test (*76*) and the 5-point Bederson’s sensory-motor deficit scoring system (*77*). Briefly, the following scores were given: a 0, no motor deficit; 1, flexion of torso and contralateral forelimb when mouse was lifted by the tail; 2, circling to the contralateral side when mouse is held by the tail on a flat surface, but normal posture at rest; 3, leaning to the contralateral side at rest, 4, no spontaneous motor activity, 5, early death due to stroke.

Infarct size was calculated based on cresyl-violet stained coronal sections as described previously in (*37, 78, 79*). In brief, lesion volume at 24 h reperfusion was calculated by integration of areas of damage measured at eight neuro-anatomically defined coronal levels (between 2.9 mm rostral and 4.9 mm caudal to bregma) followed by correction for oedema.

To delineate the ischemic penumbra, unfixed 1 mm thick brain slices were incubated in 1% TTC (2,3,5-Triphenyltetrazolium chloride, Sigma) dissolved in PBS at 37°C for 20 minutes. Slices were then postfixed with 4% PFA in PB for 24 hours at 4°C, resectioned and processed for immunostaining.

### *In vivo* two-photon imaging

Animals were anaesthetized using fentanyl. Cranial window (3 mm diameter) was opened on the left hemisphere above the primary somatosensory area and supplementary somatosensory area border (centered 3 mm lateral and 2 mm posterior to bregma) without hurting the dura mater. After removal of the skull bone a 3 mm and 5 mm double glass coverslip contruct was fixed with wetbond tissue glue on top of the dura mater. Then a custom made metal headpiece (Femtonics Ltd., Budapest, Hungary) was fixed with dental cement on the surface of the skull. All experiments were performed on a Femto2D-DualScanhead microscope (Femtonics Ltd., Budapest, Hungary) coupled with a Chameleon Discovery laser (Coherent, Santa Clara, USA). For tDTomato electroporated animals the wavelength of the laser was set to 920 nm to measure the tdTomato and GFP signals simultaneously. For Mito-R-GECO1 mitochondrial electroporated animals the wavelength was set to 1000 nm. Following excitation the fluorescent signal was collected using a Nikon 18x water immersion objective. Data acquisition was performed by MES software (Femtonics Ltd.). Since it has recently been shown that volatile anesthetics such as isoflurane may influence microglial process motility (*61*), we used fentanyl anaesthesia for these studies, which did not block microglial responses (Fig. S3c).

To analyze contacts established by microglial processes on neuronal cell bodies and proximal dendrites, we used *in utero* tdTomato electroporated CX3CR1^+/GFP^ mice. To visualize microglial processes and neuronal mitochondria simultaneously, we used Mito-R-GECO1 electroporated CX3CR1^+/GFP^ mice. Galvano Z-stacks of 7 images (820×820 pixels, 5 µm step size, range=200-225 µm from pial surface) were made at every 2 or 2.5 minutes. Two-photon image sequences were exported from MES and analyzed using FIJI. Dual colour images were analyzed with the Manual Tracking plugin of FIJI. We applied a local maximum centring correction method with a search square of 5 pixels. Pixel size was 167 nm/px.

Microglial process velocity was measured on time-series images acquired with 2P microscopy. Following motion correction, monocolour images from the same region of CX3CR1^+/GFP^ mice taken 135 seconds apart were analyzed with the Manual Tracking plugin of FIJI. We applied a local maximum centring correction method with a search square of 5 pixels. Pixel size was 167 nm/px. Processes were included in the measurement, when they were clearly traceable for at least 10 minutes.

The GCaMPf signal was imaged with the laser wavelength set to 920 nm, using the resonant scanner at 32.75 Hz. Image size was 512×488 pixels.

### Immunofluorescent labeling and confocal laser-scanning microscopy (CLSM)

Before the immunofluorescent staining, the 50 µm thick sections were washed in PB and Tris-buffered saline (TBS). This was followed by blocking for 1 hour in 1% human serum albumin (HSA) and 0.1% Triton X-100 dissolved in TBS. After this, sections were incubated in mixtures of primary antibodies overnight at room temperature. After incubation, sections were washed in TBS, and were incubated overnight at 4°C in the mixture of, all diluted in TBS. Secondary antibody incubation was followed by washes in TBS, PB, the sections were mounted on glass slides, and coverslipped with Aqua-Poly/Mount (Polysciences). Immunofluorescence was analyzed using a Nikon Eclipse Ti-E inverted microscope (Nikon Instruments Europe B.V., Amsterdam, The Netherlands), with a CFI Plan Apochromat VC 60XH oil immersion objective (numerical aperture: 1.4) and an A1R laser confocal system. We used 405, 488, 561 and 647 nm lasers (CVI Melles Griot), and scanning was done in line serial mode. Image stacks were obtained with NIS-Elements AR software, and deconvolved using Huygens Professional software (www.svi.nl). For primary and secondary antibodies used in this study, please see Supplementary Table 3.

### Quantitative analysis of CLSM data

Quantitative analysis of each dataset was performed by at least two observers, who were blinded to the origin of the samples, the experiments and did not know of each other’s results. For the analysis of somatic junction prevalence, confocal stacks with double immunofluorescent labeling (cell type-marker and microglia) were acquired from at least three different regions of mouse cortex. All labeled and identified cells were counted, when the whole cell body was located within the Z-stack. A somata was considered to be contacted by microglia, when a microglial process clearly touched it.

For the analysis of synaptic contact prevalence, confocal stacks with triple immunofluorescent labeling (pre- and postsynaptic markers and microglia) were analyzed in an unbiased, semi-automatic method. First, the two channels representing the pre- and postsynaptic markers were exported from a single image plane. The channels were thresholded automatically in FIJI, the „fill in holes” and „erode” binary processes applied. After automatic particle tracking, synapses were identified where presynaptic puncta touched postsynaptic ones. From these identified points we selected 200/animal in a systematic random manner. After this, the corresponding synapses were found again in the original Z-stacks. A synapse was considered to be contacted by microglia, when a microglial process was closer than 200 nm (4 pixels on the images).

To measure the distribution of Kv2.1 labeling relative to microglial processes, confocal stacks were exported into single-channel TIFF-series. Identical measuring frames (1,32 µm^2^) were placed randomly along the surface of pyramidal cells and integrated Kv2.1 fluorescent density was measured in each frame in FIJI. Afterwards, frames containing microglial contacts were identified (“contact” group) and compared with frames not containing microglial processes (“non-contact” group).

For the measurements of mitochondrial fragmentation we used tissue from mice that were sacrificed 4 hours after a one hour long unilateral MCAo. Confocal stacks with double immunofluorescent labeling (Kv2.1 and TOM20) were taken from the penumbra and the corresponding contralateral region. We used the Kv2.1 labeling to trace neuronal cell bodies as regions of interest (ROI). Every cell was counted once using the confocal plane containing its largest cross-section. Within the ROIs TOM20 labeling was investigated with FIJI: after automatic thresholding we ran the ‘Analyze Particles’ command to obtain the area and the major axis of individual somatic mitochondria. For the analysis of Kv2.1 clusters, individual cells were measured by using the confocal Z-plane containing the largest cross-section of the cell body. The intensity profile of Kv2.1 labeling was plotted using FIJI. A cluster was identified when at least three adjacent pixels’ intensity was more than 25 grayscale values (10% of an 8-bit image) larger than the average fluorescent intensity of that particular cells Kv2.1 labeling.

Microglial process coverage was measured on CLSM Z-stacks acquired with a step size of 300 nm. On single-channel images, Kv2.1-positive cells were selected randomly, the cell bodies of which were fully included in the captured volume. The surface of these cells was calculated by measuring the circumference of the soma on every section multiplied by section thickness. The length of microglial process contacts was measured likewise.

TOM20 and vesicular nucleotide transporter (vNUT) fluorescent intensity profiles were analysed using a semi-automatic method. Confocal stacks with triple immunofluorescent labeling (microglia, Kv2.1 and TOM20/vNUT) were collected. The section containing the largest cross-section of a pyramidal cell was used to trace the cell membrane according to Kv2.1-labeling. This contour was then expanded and narrowed by 0.5 µm to get an extracellular and an intracellular line, respectively. The intensity of fluorescent labeling was analyzed along these lines. After normalizing and scaling, microglial contact was identified where microglial fluorescent intensity was over 20% of total, for at least 500 nm. Then the contact area was extended 500-500 nm on both sides, and TOM20/VNUT fluorescent intensity within these areas were measured for “contact” value.

3-dimensional reconstruction of CLSM and 2P imaging stacks was performed using the IMOD software package (*80*).

### STORM superresolution imaging

Free-floating brain sections were blocked with 2% normal donkey serum followed by immunostaining with rabbit anti-P2Y12R and mouse anti-Kv2.1 antibodies, followed by anti-rabbit Alexa 647 and anti-mouse Alexa 594 secondary antibodies. Sections were mounted onto #1.5 borosilicate coverslips and covered with imaging medium containing 5% glucose, 0.1 M mercaptoethylamine, 1 mg/ml glucose oxidase, and catalase (Sigma, 1500 U/ml) in Dulbecco’s PBS (Sigma), immediately before imaging (*81*). STORM imaging was performed for P2Y12R (stimulated by a 647 nm laser) by using a Nikon N-STORM C2+ superresolution system that combines ‘Stochastic Optical Reconstruction Microscopy’ technology and Nikon’s Eclipse Ti research inverted microscope to reach a lateral resolution of 20 nm and axial resolution of 50 nm (*23, 82*). Imaging was performed using the NIS-Elements AR 4.3 with N-STORM 3.4 software, and we used VividSTORM open source software (*23*). Molecule lists were exported from NIS in txt format, and the three image planes of the ics-ids file pairs from the deconvolved confocal stacks matching the STORM volume were converted to the ome-tiff format using Fiji software. Confocal and corresponding STORM images were fitted in VividSTORM. Localization points exceeding a photon count of 2000 were counted as specific superresolution localization points. Local density filter (10 neighbours within 150 nm for P2Y12R and IBA1, and 5 neighbours within 150 nm for Kv2.1) and Z-filter (±300 nm from focal plane) was applied to the localization points.

### Pre-embedding immunoelectron microscopy

After extensive washes in PB and 0.05 M Tris-buffered saline (pH 7.4; TBS) sections were blocked in 1 % human serum albumin (HSA; Sigma-Aldrich) in TBS. Then, they were incubated in primary antibodies (Supplementary Table 3) diluted in TBS containing 0.05% sodium azide for 2-3 days. After repeated washes in TBS, the sections were incubated in blocking solution (Gel-BS) containing 0.2 % cold water fish skin gelatine and 0.5 % HSA in TBS for 1 h. Next, sections were incubated in gold-conjugated or biotinylated secondary antibodies (Supplementary Table 3) diluted in Gel-BS overnight. After extensive washes in TBS the sections were treated with 2 % glutaraldehyde in 0.1 M PB for 15 min to fix the gold particles into the tissue. This was occasionally followed by incubation in avidin– biotinylated horseradish peroxidase complex (Elite ABC; 1:300; Vector Laboratories) diluted in TBS for 3 h at room temperature or overnight at 4°C. The immunoperoxidase reaction was developed using 3,3-diaminobenzidine (DAB; Sigma-Aldrich) as chromogen. To enlarge immunogold particles, sections were incubated in silver enhancement solution (SE-EM; Aurion) for 40-60 min at room temperature. The sections were then treated with 1% (for electron tomography) or 0.5 % OsO4 in 0.1 M PB, at room temperature (for electron tomography) or on ice, dehydrated in ascending alcohol series and in acetonitrile and embedded in Durcupan (ACM; Fluka). During dehydration, the sections were treated with 1% uranyl acetate in 70% ethanol for 20 min. For electron microscopic analysis, tissue samples from the CA1 area of dorsal hippocampus/somatosensory cortex (S1) were glued onto Durcupan blocks. Consecutive 70 nm thick (for conventional electron microscopic analysis) or 150 nm thick (for electron tomography) sections were cut using an ultramicrotome (Leica EM UC6) and picked up on Formvar-coated single-slot grids. Ultrathin sections for conventional electron microscopic analysis were and examined in a Hitachi 7100 electron microscope equipped with a Veleta CCD camera (Olympus Soft Imaging Solutions, Germany). 150 nm thick electron tomography sections were examined in FEI Tecnai Spirit G2 BioTwin TEM equipped with an Eagle 4k HS camera.

### Electron tomography

For the electron tomographic investigation we used 150 nm thick sections from the hippocampal CA1 region from the anti-P2Y12R immunogold stained material (see: “Pre-embedding immunoelectron-microscopy”). Before electron tomography, serial sections on single-slot copper grids were photographed with a Hitachi H-7100 electron microscope and a Veleta CCD camera. After this, grids were put on drops of 10% HSA in TBS for 10 minutes, dipped in distilled water (DW), put on drops of 10 nm gold conjugated Protein-A (Cytodiagnostics #AC-10-05) in DW (1:3), and washed in DW. Finally, we deposited 5 nm thick layers of carbon on both sides of the grids. Electron tomography was performed using a Tecnai T12 BioTwin electron microscope equipped with a computer-controlled precision stage (CompuStage, FEI). Acquisition was controlled via the Xplore3D software (FEI). Regions of interest were pre-illuminated for 4-6 minutes to prevent further shrinkage. Dual-axis tilt series were collected at 2 degree increment steps between -65 and +65 degrees at 120 kV acceleration voltage and 23000× magnification with -1.6 – -2 µm objective lens defocus. Reconstruction was performed using the IMOD software package (*80*). Isotropic voxel size was 0.49 nm in the reconstructed volumes. After combining the reconstructed tomograms from the two axes, the nonlinear anisotropic diffusion filtering algorithm was applied to the volumes. Segmentation of different profiles has been performed on the virtual sections using the 3Dmod software, and measurements were done on the scaled 3D models.

Analysis of the connection between membrane distance and P2Y12R density was carried out by investigating all points of the microglial membrane facing neuronal soma, using reconstructed 3D models. Coordinates of the points of neuronal membrane, soma-facing microglial membrane and P2Y12R labeling gold particles were exported using IMOD. For every single point of microglial membrane, the lowest distance to the neuronal membrane and the number of P2Y12R labeling gold particles within 40 nm was calculated with a unique algorithm running in program R (The R Foundation). Since neuronal junctions established by microglial processes are dynamic, a strong linear correlation can not be expected, therefore statistical analysis was carried out by dividing data into two groups. In the analysed tomograms the average distance between neuronal somatic membranes and facing microglial membranes was 13.06 nm, which we used as demarcation point.

Analysis of P2Y12R density along different surfaces of microglial processes was done using reconstructed 3D models. We identified segments of microglial membranes facing (running parallel with) different neuronal membranes. These segments were grouped depending on opposing neuronal profiles (e.g neuronal soma or other neuronal parts). Using IMOD, the surfaces of microglial profiles were measured and gold particles were assigned to the closest membrane part. Due to different labeling density and penetration differences, we only performed pairwise comparisons between ‘somatic’ and ‘non-somatic’ microglial membranes within the same microglial processes. Only those gold particles were counted that localized within 40 nm of the microglial membrane.

### *In vitro* nicotinamide adenine dinucleotide (NADH) imaging

Mice were anaesthetized by inhalation of halothane, and following decapitation 200 µm thick coronal slices were prepared from the somatosensory and visual cortex with a vibrating blade microtome (Microm HM 650 V) immersed in slicing solution containing (in mM): 130 NaCl, 3.5 KCl, 1 NaH2PO4, 24 NaHCO3, 1 CaCl2, 3 MgSO4, 10 D(+)-glucose, saturated with 95% O2 and 5% CO2. The solution used during experiments was identical to the slicing solution, except it contained 3 mM CaCl2 and 1.5 mM MgSO4. Experiments were carried out less than 4 hours after slicing. During image acquisition slices were kept at ∼36°C. Imaging with multiphoton excitation was performed using a Zeiss LSM 7MP scanning microscope (Carl Zeiss, Germany) through a 40× water-immersion objective (W-Plan, NA 1.0, Carl Zeiss). To acquire simultaneous excitation of GFP and NADH autofluorescence of acute brain slices we used two single wavelength mode-locked Ti:sapphire lasers the beams of which were coupled to each other by a dichroic beam splitter (t810lpxr, Chroma Technology Corp, USA). One of our lasers (MaiTai DeepSee, Spectra-Physics, Santa Clara, USA) exciting GFP operated at 885 nm, while our second laser (FemtoRose 100 TUN, R&D Ultrafast Lasers, Hungary) exciting NADH had an operation wavelength of 750 nm. Both laser systems delivered ∼100 fs pulses at ∼80 MHz and ∼76 MHz repetition rate, respectively. A total average laser power of 16-18 mW was measured after the objective during imaging. Time-lapse images (1024×1024 pixels) were collected continuously for up to 55 minutes with 30.98 s frame scan time. Emission filters were chosen to separate intrinsic NADH fluorescence (ET460/50m, Chroma Technology Corp, USA) from GFP fluorescence (ET525/50m, Chroma Technology Corp, USA). Time lapse images were processed and analyzed in Fiji (ImageJ, NIH) software. As a first step images were spatial filtered (mean filter smooth with 1 pixel diameter) and corrected for contrast. To remove jitter in image series stabilization was applied with a FIJI plugin (K. Li, “The image stabilizer plugin for ImageJ,” http://www.cs.cmu.edu/ ∼kangli/code/Image_Stabilizer.html, February, 2008). Analysis and quantification of NADH fluorescence was carried out in manually selected areas of compartments of cell bodies at microglial contact site.

### *In vivo* widefield calcium imaging

*In vivo* widefield calcium imaging was performed as previously described in detail (*83*). In brief, as an optogenetic calcium-reporter mouse strain, C57BL/6J-Tg(Thy1-GCaMP6s)GP4.12Dkim/J (*72*) heterozygous mice were bred at the Institute for Stroke and Dementia Research, Munich. The skin covering the skull and the underlying connective tissue were removed in head-fixed mice and a layer of transparent dental cement was distributed on the window area and covered with a coverslip. Afterwards, the mice were allowed to recover from the surgery for more than 48 h before the first image acquisition. For image acquisition, mice were injected with 0.05 mg/kg bodyweight of medetomidine intraperitonially 5 minutes prior to inducing inhalation anesthesia with a mixture of 5% isoflurane in 70% nitrous oxide and 30% oxygen. After 70 seconds, the animals were fixed in a stereotactic frame, the dose of isoflurane was decreased to 1.5% for 140 seconds and finally decreased to 0.75% for 4 minutes to maintain steady-state before data-acquisition. *In vivo* widefield calcium imaging was performed on a custom-built imaging setup described in (*83*). This setup allowed widefield imaging through the chronic window on top of the skull into the cortex of both forebrain hemispheres by covering a field-of-view of 12×12 mm, corresponding to an image matrix of 330×330 pixels. Image acquisition was conducted for 29 minutes (44 × 1000 frames, immediately after MCAo induction) or 4 minutes (6 × 1000 frames, baseline acquisition, 60 min after injection of PSB0739 and after 120 min reperfusion, respectively). After the imaging session, anesthesia was terminated by intraperitoneally injecting the mice with 0.1 mg/kg bodyweight Atipamezole. During all anesthetized procedures body temperature was maintained using a feedback-controlled heating system. After end of surgeries animals were put in a heating chamber until they had recovered from anesthesia. Post-surgery analgesia and sedation protocols were conducted in accordance with approved protocols by the governmental committee.

Data processing was performed in MATLAB (R2016b, The MathWorks, USA). Briefly, all calcium images were first resized to 2/3 to a uniform image matrix of 220×220 pixels and resolution of 18.3 pixel/mm, followed by a normalization of the fluorescence signal (ΔF/F) and bandpass filtering (0.1 to 1 Hz). From the filtered signal time courses 10 seconds were cropped from beginning and end to reduce filter artifacts. From the preprocessed data files, videos were created to evaluate recording and preprocessing quality. The videos were reviewed by an experienced rater and excluded when containing motion artifacts. To ensure intra- and interindividual correspondence of the analyzed cortical regions, preprocessed images from different acquisition blocks and animals were spatially co-registered, as previously described (*83*). In the registered images, the bregma was always located in the center of the image matrix and the sagittal suture corresponded to a vertical line.

Data processing was performed in MATLAB (R2016b, The MathWorks, USA). Briefly, all calcium images were first resized to 2/3 to a uniform image matrix of 220×220 pixels and resolution of 18.3 pixel/mm, followed by a normalization of the fluorescence signal (ΔF/F) and bandpass filtering (0.1 to 1 Hz). From the filtered signal time courses 10 seconds were cropped from beginning and end to reduce filter artifacts. From the preprocessed data files, videos were created to evaluate recording and preprocessing quality. The videos were reviewed by an experienced rater and excluded when containing motion artifacts. To ensure intra- and interindividual correspondence of the analyzed cortical regions, preprocessed images from different acquisition blocks and animals were spatially co-registered, as previously described (*83*). In the registered images, the bregma was always located in the center of the image matrix and the sagittal suture corresponded to a vertical line.

Images of every acquisition were masked in a two-step procedure. First, a general mask was applied to exclude lateral cortical areas, which were out of focus due to the curvature of the cortical surface as described in (*83*). Second, an individual mask was computed to exclude all pixels, in which the calcium signal was saturated due to autofluorescence, as occurring in areas affected by the infarct (*83*). Both, the general and the individual masks were combined for every acquisition.

To characterize changes in the cortical network after stroke, functional connectivity was computed between pairs of ROIs, representing functional cortical areas, previously defined (*83*). Functional connectivity was calculated as the Fisher z-transformed Pearson-moment correlation between the ROI signal time-courses. The average connectivity scores were calculated within each group (PSB0739 treated and control group), and the difference between groups were depicted for all ROI pairs in a heatmap.

For seed-based functional connectivity analysis, connectivity scores were calculated in the same way but between a selected ROI in the right hindlimb sensory area (rHLs) and the signal time series of all pixels on the cortical surface included in the combined general and individual masks. In order to quantify connectivity change of the rHLs after stroke, the connectivity scores were normalized by deviding through the connectivity scores resulting in the baseline condition. Results were visualized as topographical maps of all brain pixel.

Overall functional connectivity alterations due to treatment of the mice with PSB0739 (naïve and stroke) were assessed by computing the global connectivity (*84, 85*) for each pixel inside the combined general and individual mask. For a given pixel, global connectivity was calculated by calculating the functional connectivity with each other pixel inside the mask, followed by averaging across the resulting connectivity scores. In order to assess treatment effects, global connectivity scores were averaged pixel-wise within group (PSB0739 and control group). The difference between groups was than visualized as topographical map of all brain pixels.

To compare the extent of global connectivity dropdown between PSB0739 and control treated animals after stroke, the sum of all pixels with a moderate global connectivity (i.e. global connectivity less than 0.6 according to (*86*)) was calculated for every mouse. The mean area of each group was contoured within the global connectivity map and the area was assessed quantitatively per animal and compared between the groups.

To represent the non-functionality of cortical tissue during the occlusion of the middle cerebral artery, the number of saturated pixels during the occlusion was calculated. During the occlusion, cortical spreading depressions (CSDs) appeared. Given the high degree of neuronal activation within areas covered by these waves, the high amplitude of the Ca^2+^ derived signal caused saturation in these areas. Therefore, we quantified the spatial extent of CSDs by counting the saturated pixels during these waves. The start- and end-time of every cortical wave were defined as first appearance of saturated pixels and the full disappearance of saturated pixels. The absolute maximum spatial extent of every cortical wave was identified and used to align the individual cortical waves of all animals. The area of saturated pixels was then acquired for every CSD wave for every animal atthe endpoint of the shortest wave (which ended 37 seconds after the aligned absolute maximum). The area of saturated pixels was groupwise depicted as overlay of the area of every CSD wave upon the general mask.

### Statistical analysis

All quantitative assessment was performed in a blinded manner and based on power calculation wherever it was possible. Based on the type and distribution of data populations (examined with Shapiro-Wilks W test) we applied appropriate statistical tests: in case of two independent groups of data unpaired t-test or Mann Whitney U-test, for two dependent groups of data Wilcoxon signed-rank test, for multiple comparisons one-way ANOVA (with Tukey’s test) or Kruskal-Wallis test was used (indicated accordingly in the main text). Statistical analysis was performed with the Statistica 13.4.0.14 package (TIBCO), differences with p<0.05 were considered significant throughout this study.

## Supplementary Material

**Table S1.**

| Localisation point density (LP/um2) |  |  |  |  |  |  |
| --- | --- | --- | --- | --- | --- | --- |
| animal | category | mean | S.E.M. | n | comparison (Tukey's multiple comparisons test) | p-value |
| GCamp6-injected | contact | 136.8 | 10.02 | 12 | contact vs. non-contact | <0,0001 |
|  | non-contact | 60.42 | 7.719 | 12 | contact vs. whole microglia | <0,0001 |
|  | whole microglia | 76.8 | 5.128 | 12 | whole microglia vs. non-contact | 0.3179 |
| camK2A/gfp/22 | contact | 142.6 | 17.17 | 18 | contact vs. non-contact | <0,0001 |
|  | non-contact | 52.2 | 10.08 | 17 | contact vs. whole microglia | 0.0008 |
|  | whole microglia | 76.47 | 6.054 | 18 | whole microglia vs. non-contact | 0.3423 |
| gad65_3e/gfp 5.5/30 | contact | 81.05 | 15.37 | 11 | contact vs. non-contact | 0.0116 |
|  | non-contact | 38.62 | 4.892 | 13 | contact vs. whole microglia | 0.0197 |
|  | whole microglia | 41.58 | 7.661 | 13 | whole microglia vs. non-contact | 0.973 |
| BAC_pva/gfp/2 | contact | 93.75 | 21.59 | 10 | contact vs. non-contact | 0.0009 |
|  | non-contact | 20.03 | 2.746 | 10 | contact vs. whole microglia | 0.003 |
|  | whole microglia | 31.17 | 5.736 | 12 | whole microglia vs. non-contact | 0.7957 |
| Cluster density (cluster/um2) |  |  |  |  |  |  |
| animal | category | mean | S.E.M. | n | comparison (Tukey's multiple comparisons test) | p-value |
| GCamp6-injected | contact | 0.9705 | 0.084 | 12 | contact vs. non-contact | <0,0001 |
|  | non-contact | 0.4451 | 0.071 | 12 | contact vs. whole microglia | 0.0004 |
|  | whole microglia | 0.5401 | 0.055 | 12 | whole microglia vs. non-contact | 0.6164 |
| camK2A/gfp/22 | contact | 0.6089 | 0.069 | 18 | contact vs. non-contact | 0.0002 |
|  | non-contact | 0.3111 | 0.046 | 18 | contact vs. whole microglia | 0.0031 |
|  | whole microglia | 0.3673 | 0.02 | 18 | whole microglia vs. non-contact | 0.7007 |
| gad65_3e/gfp 5.5/30 | contact | 0.368 | 0.072 | 9 | contact vs. non-contact | 0.4608 |
|  | non-contact | 0.2659 | 0.052 | 13 | contact vs. whole microglia | 0.4681 |
|  | whole microglia | 0.2684 | 0.051 | 14 | whole microglia vs. non-contact | 0.9994 |
| BAC_pva/gfp/2 | contact | 0.3726 | 0.062 | 10 | contact vs. non-contact | 0.0107 |
|  | non-contact | 0.1839 | 0.024 | 11 | contact vs. whole microglia | 0.0553 |
|  | whole microglia | 0.2293 | 0.036 | 12 | whole microglia vs. non-contact | 0.7145 |

**Table S2.**

| Subject | Code | Gender | Age (years) | Health status | Survival after stroke (days) | Comorbidities | Cause of death | Tissue sample type |
| --- | --- | --- | --- | --- | --- | --- | --- | --- |
| Stroke patient | 2011/0092 | female | 77 | normal | 2 | arterial hypertension, type II diabetes, hyperthyreosis | stroke | paraffin embedded sections |
| Stroke patient | 2003/0029 | male | 66 | preceeding systemic inflammatory burden | 1 | arterial hypertension | stroke | paraffin embedded sections |
| Stroke patient | 2014/0050 | female | 78 | normal | 1 | unknown | stroke | paraffin embedded sections |
| Control subject | SKO3 | female | 59 | normal | n.a. | ischemic cardiomyopathy | cardiogenic shock | free floating and paraffin sections |
| Control subject | SKO13 | female | 60 | normal | n.a. | chronic bronchitis | respiratory arrest | free floating and paraffin sections |
| Control subject | SKO16 | male | 73 | normal | n.a. | atherosclerosis, pneumonia | respiratory arrest | free floating and paraffin sections |

**Table S3.**

| Primary antibodies | host | source | catalog nr. | RRID |
| --- | --- | --- | --- | --- |
| CB1 | goat | MyBioSource | MBS422809 | n.a. |
| CytC | mouse | BioLegend | 612302 | RRID:AB_315774 |
| c-fos | guinea-pig | Synaptic Systems | 226 004 | RRID:AB_2619946 |
| Gephyrin | mouse | Synaptic Systems | 147 021 | RRID:AB_2232546 |
| GFP | chicken | Invitrogen | A10262 | RRID:AB_2534023 |
| Homer1 | chicken | Synaptic Systems | 160 006 | RRID:AB_2631222 |
| Iba1 | goat | Novusbio | NB100-1028 | RRID:AB_521594 |
| Iba1 | rabbit | Wako Chemicals | 019-19741 | RRID:AB_839504 |
| Kv2.1 | mouse | NeuroMab | 75-014 | RRID:AB_10673392 |
| Kv2.1 | rabbit | Synaptic Systems | 231 002 | RRID:AB_2131650 |
| MAP2 | guniea-pig | Synaptic Systems | 188 004 | RRID:AB_2138181 |
| P2Y12R | rabbit | Anaspec | AS-55042A | RRID:AB_2267540 |
| P2Y12R | rabbit | Anaspec | AS-55043A | RRID:AB_2298886 |
| Parvalbumin | goat | Swant | PVG 213 | RRID:AB_2721207 |
| RFP | rat | ChromoTek | 5f8-100 | RRID:AB_2336064 |
| SMI32 | mouse | Covance | SMI-32P | RRID:AB_10719742 |
| TOM20 | rabbit | Santa Cruz | sc-11415 | RRID:AB_2207533 |
| vGAT | guinea pig | Synaptic Systems | 131 004 | RRID:AB_887873 |
| vGluT1 | guinea pig | Millipore | AB5905 | RRID:AB_2301751 |
| vGluT3 | guinea pig | Synaptic Systems | 135 204 | RRID:AB_2619825 |
| VNUT | guinea pig | Millipore | ABN83 | n.a. |
| Secondary antibodies |  |  |  |  |
| biotinylated anti-rabbit | donkey | BioRad | 644008 | RRID:AB_619842 |
| DyLight 405 anti-mouse | donkey | Jackson | 715-475-150 | RRID:AB_2340839 |
| Alexa 405 anti-rabbit | donkey | Jackson | 711-475-152 | RRID:AB_2340616 |
| Alexa 488 anti-chicken | donkey | Jackson | 703-546-155 | RRID:AB_2340376 |
| Alexa 488 anti-goat | donkey | Jackson | 705-546-147 | RRID:AB_2340430 |
| Alexa 488 anti-rabbit | donkey | Jackson | 711-546-152 | RRID:AB_2340619 |
| CF568 anti-mouse | donkey | Biotium | 20802 | RRID:AB_10853136 |
| Alexa 594 anti-goat | donkey | LifeTech | A11058 | RRID:AB_2534105 |
| Alexa 594 anti-guinea pig | goat | LifeTech | A11076 | RRID:AB_141930 |
| Alexa 594 anti-mouse | donkey | LifeTech | A21203 | RRID:AB_141633 |
| Alexa 594 anti-rabbit | donkey | LifeTech | A21207 | RRID:AB_141637 |
| Alexa 594 anti-rat | donkey | Jackson | 712-585-150 | RRID:AB_2340688 |
| Alexa 647 anti-goat | donkey | Jackson | 705-606-147 | RRID:AB_2340438 |
| Alexa 647 anti-guinea pig | donkey | Jackson | 706-606-148 | RRID:AB_2340477 |
| Alexa 647 anti-mouse | donkey | Jackson | 715-605-150 | RRID:AB_2340866 |
| Alexa 647 anti-rabbit | donkey | Jackson | 711-605-152 | RRID:AB_2492288 |

## Supplementary Figures

**Figure S1.**
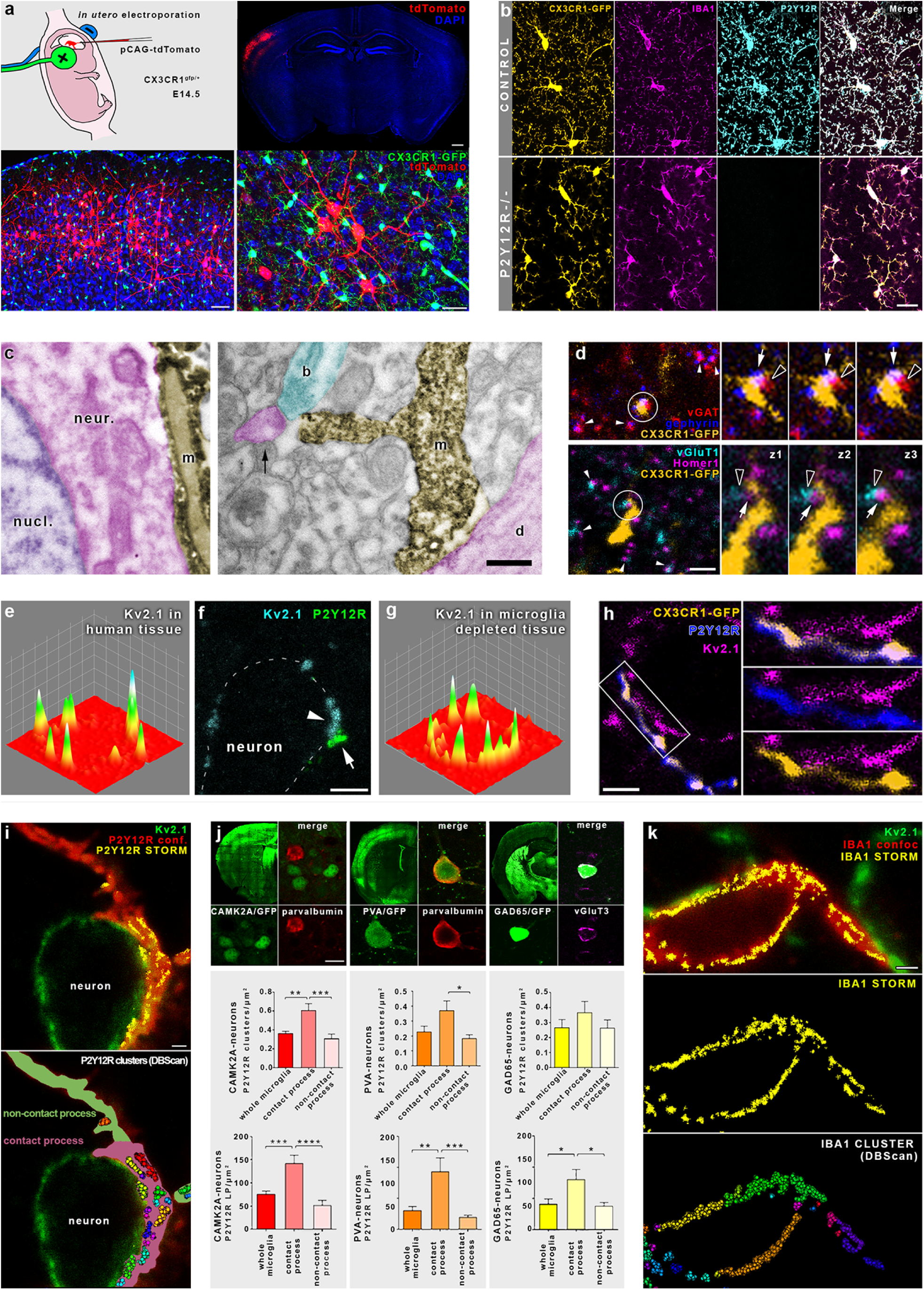
a) Schematic illustration and result of CX3CR1+/GFP mouse in utero electroporation with pCAG-IRES-tdTomato. Upper right shows the outspread of tdTomato-positive neurons in red. Lower panels show the non-overlapping staining of electroporated neurons (red) and microglia (green). Cell nuclei are visible in blue. b) CLSM images showing three microglial markers. Upper row shows complete overlap of immunolabelings against CX3CR1 (yellow), Iba1 (magenta) and P2Y12R (cyan) in CX3CR1+/GFP mice. Lower row shows the presence of overlapping CX3CR1 and Iba1 signals with the complete absence of P2Y12R labeling in P2Y12R-/- animals. c) Electron microscopic images showing microglial processes (m) establishing direct contact with neuronal cell body (left, n), dendritic shaft (d) and presynaptic bouton (b) in mice. Microglia are visualized by immunoperoxidase reaction against Iba1. Pseudocoloring shows microglia in yellow, neuronal cell body and dendritic processes in magenta, presynaptic bouton in cyan and nucleus (nucl.) in purple. Arrow points at a dendritic spine receiving asymmetric synaptic contact. d) Microglial processes contacting cortical inhibitory and excitatory synapses. Presynaptic terminals are visible by stainings against vGAT (red) and vGluT1 (cyan), while postsynaptic side is characterized by stainings against gephyrin (blue) and Homer1 (magenta). White arrowheads point at colocalization of pre- and postsynaptic markers. Microglial contacts in white circles are enlarged in the panels right, with the previous and following Z-planes from the stack (Z-step = 250 nm). Empty arrowheads point at the presynaptic marker, white arrows point at the postsynaptic side. e) Heatmap shows Kv2.1 clusters of a human cortical neuron. f) CLSM image shows that a P2Y12R-labeled microglial process contacts a human cortical neuron exactly at the spot where a strong Kv2.1 cluster is located. g) Heatmap shows that Kv2.1-clustering of mouse cortical neurons is not affected by microglia depletion. i) P2Y12R clustering differs depending on location. Microglial processes were classified depending on neuronal contacts established: contact processes are pale green, non-contact processes are pale magenta on bottom panel. j) Analysis of P2Y12R cluster and LP density on different microglial processes contacting distinct cell populations. Microglial contact processes on CamK2a (red) and PVA-positive cells (orange) possess significantly higher cluster density, than non-contact segments, while there is a similar trend observable in the case of GAD65-positive cells (yellow, for detailed results see Supplementary Table 1.). Contact processes had significantly higer number of P2Y12R LP-s in all populations. Anti-PV and vGluT3 labelings were used to confirm cell specificity. k) Superresolution imaging of microglial Iba1 shows no clustering at somatic junctions. CLSM images show neuronal Kv2.1 (green) and a contacting microglial process made visible by Iba1-labeling (red). Iba1 (yellow) STORM-signal is overlaid and shown individually in the middle panel. Lower panel shows a homogenous distribution of Iba1 LPs using DBScan analysis. Scale bars: 500 µm on top right panel of a, 50 µm on bottom left panel of a, 25 µm on bottom right panel of a, 20 µm on b, 1 µm on c, i and k, 2 µm on d and f, 10 µm on j.

**Figure S2.**
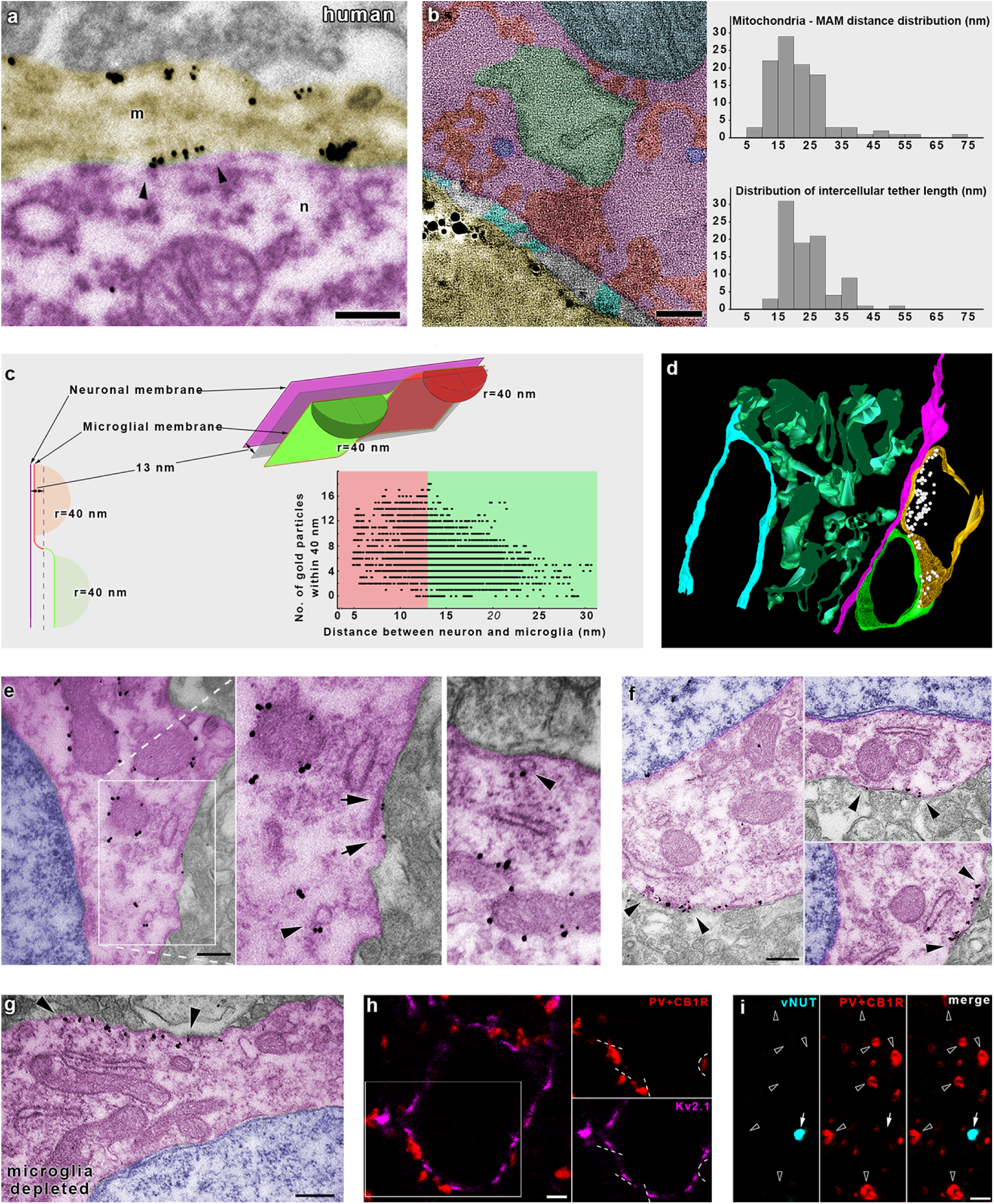
a) Electron microscopic image shows a microglial process (m) establishing direct contact with neuronal cell body (n) in human tissue. Microglia is visualized by immunogold-labeling against P2Y12R. Arrowheads point at accumulation of gold particles at contact sites. b) Single virtual plane of an electron tomographic volume (thickness: 0.49 nm) showing a P2Y12R-positive (gold particles) microglial process (yellow) contacting a neuronal cell body (magenta). Mitochondria (blue), mitochondria-associated membranes (MAM, green), vesicles (purple) and intracellular densities (red) are commonly visible in the vicinity of these junctions. In some cases, intercellular linking structures (cyan) can be seen within the cleft. Right: distribution of measured distances between mitochondria and MAM (upper chart) and measured length of intercellular tethers (lower chart). c) Schematic illustration showing the principles of P2Y12R density and membrane distance measurements using electron tomography (Fig. 2e). Single points of the microglial membrane were divided into two groups based on their closest distance from the neuronal membrane: closer than the average distance of 13 nm (red) or farther (green). For every observed point, P2Y12R labeling gold particles were counted within a radius of 40 nm. In the bottom right corner, the distribution of measured points is shown in the case of a representative junction. d) Enlarged and rotated view of the electron tomographic model of microglia-neuron junction on Fig. 2e. e) Transmission electron micrographs show TOM20-immunogold labeling in neocortical neurons. Immunogold labeling (black grains) is specifically associated with outer mitochondrial membranes, while TOM20-positive vesicles can also be observed (arrowheads). Some immunogold particles can be found on the plasma membrane of the neurons (arrows), suggesting the exocytosis of mitochondria derived vesicles. Nucleus is blue, neuronal cytoplasm is magenta. Left and middle panel is the same as on Fig. 2h. f) Transmission electron micrographs show Kv2.1 clusters on cortical neuronal cell bodies (arrowheads). g) Transmission electron micrograph of immunogold labeled mouse tissue shows that Kv2.1-clustering and the association of the cluster with neuronal organelles is not affected by microglia depletion. h) Kv2.1 accumulation does not overlap with somatic inhibitory terminals. CLSM images show perisomatic terminals stained with antibodies against PV and CB1 (red) and neuronal Kv2.1 (magenta). Panels on the right show the area in white rectangle on the first image. Dashed lines mark clear separation between perisomatic boutons and Kv2.1-signal, n=220 perisomatic boutons tested for Kv2.1 from two mice. i) Neuronal vNUT signal (cyan) is not present in perisomatic inhibitory terminals (red), n=194 perisomatic boutons tested for vNUT from two mice. Scale bars: 500 nm on a, 50 nm on b, 300 nm on e, f and g, 2 µm on h and i.

**Figure S3.**
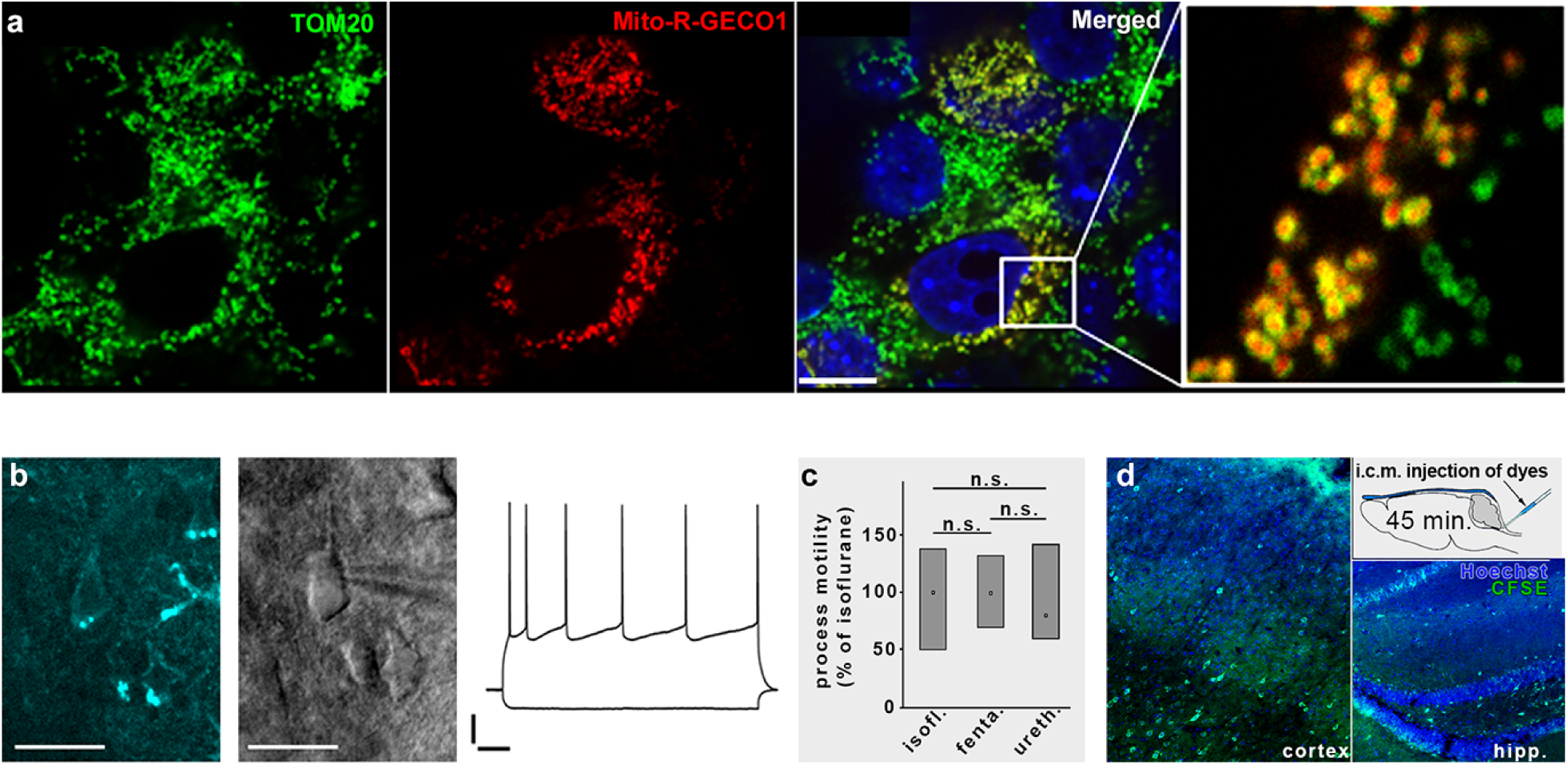
a) Neuronal cell culture transfected with CMV-Mito-R-Geco1 and counterstained with TOM20 and DAPI. Scale bar is 10 µm. b) Representative NADH intrinsic fluorescence image of a pyramidal cell. Left: Sample image of NADH fluorescence of neuronal somata. Middle: DIC image of the same region with whole cell patch clamp electrode. Right: Electrophysiological recordings of membrane potential response to -100 and +160 pA current injections. Scales: 20 mV and 100 ms. c) Effects of different anaesthetics on microglial process motility. d) Mixture of carboxyfluorescein succinimidyl ester (CFSE) and Hoechst injected into the cisterna magna (i.c.m.) rapidly diffuses to all layers of neocortex and hippocampus.

**Figure S4.**
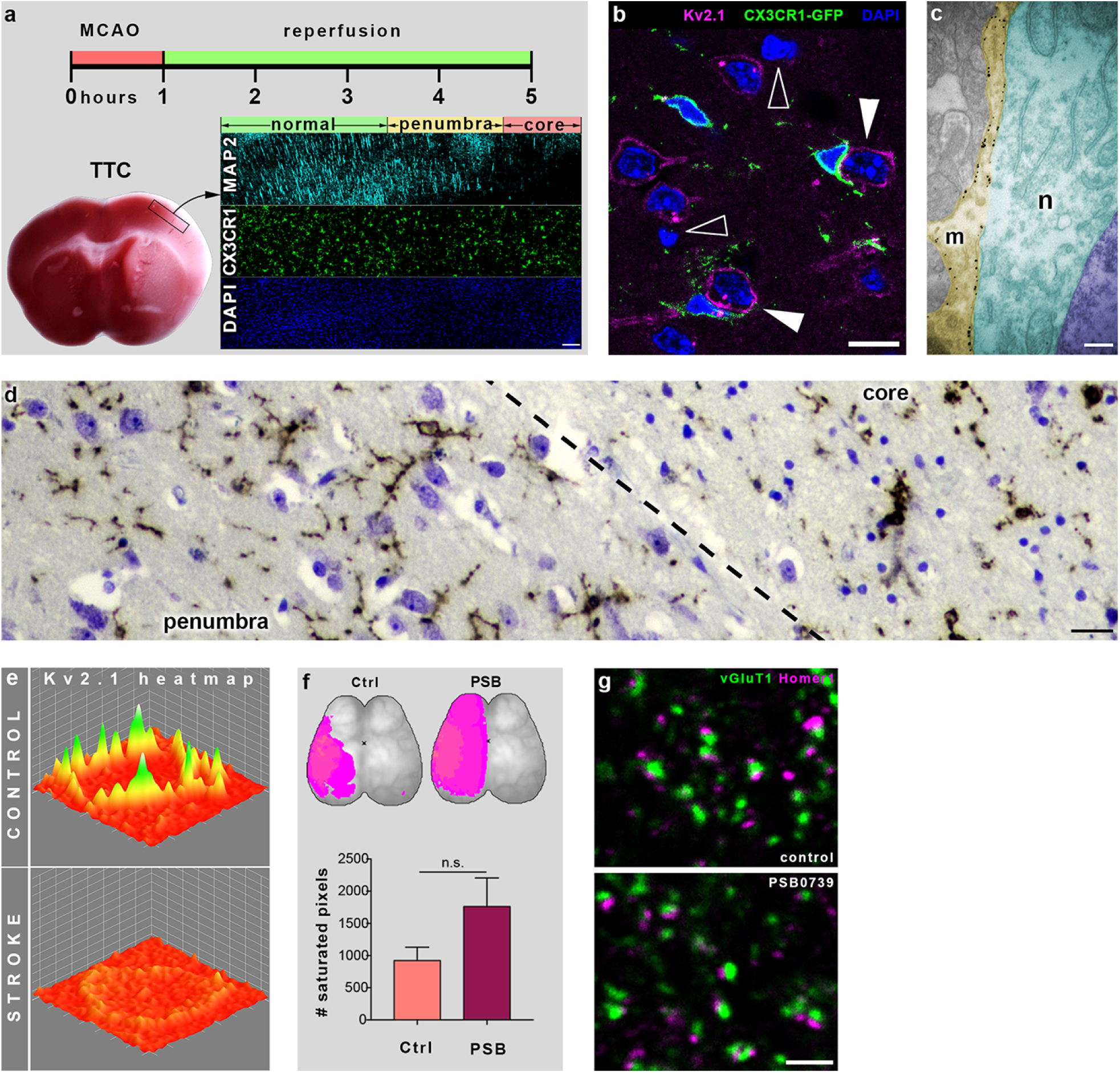
a) Outline of middle cerebral artery occlusion (MCAo) experiment and delineation of core/penumbra regions of stroke affected brain. 1 hour MCAo was followed by 4 hours of reperfusion. Delineation was performed with the use of TTC staining and immunofluorescent labeling of MAP2 and microglia. b) DAPI staining pattern reveals that neurons with increased microglial coverage (white arrowheads) in the penumbra region possess normal chromatin structure, confirming their viability. Empty arrowheads point to pycnotic nuclei. c) Transmission electron micrograph shows a P2Y12R-immunogold particle labeled microglial process covering a neuron in the penumbra region. Note the increased coverage, the disappearance of cytoplasmatic structures like closely apposed mitochondria and MAM. Neuronal membrane integrity and chromatin structure are sustained. d) Delineation of core/penumbra border in postmortem human cortical tissue. Neurons and nuclei are visualized by Nissl-staining (blue), microglia is labeled by anti-P2Y12R DAB-Ni immunoperoxidase reaction (dark brown precipitate). e) Heatmaps of Kv2.1 immunostaining reveal that Kv2.1 clusters disappear after stroke in the penumbra. f) The saturated calcium signal after experimental stroke (left MCAo in every experiment) corresponds to the dysfunctional tissue during the occlusion. The maps show an overlay of the area of saturated pixels (pink) of every animal in the two groups. g) CLSM images show type 1 vesicular glutamate transporter (vGlut1) and Homer1 immunolabeling after acute i.c.m. administration of vehicle (control) or PSB0739. The closely apposed vGluT1 and Homer1 puncta represent glutamatergic synaptic contacts. PSB0739-treatment did not alter neocortical synapse numbers. Scale bars: 100 µm on a, 10 µm on b, 300 nm on c, 12 µm on d, 2 µm on g.

## Supplementary Movie legends

***Movie S1.*** *In vivo 2P time-lapse imaging shows temporal dynamics of microglia-neuron contacts. A tdTomato expressing neocortical neuron (red) is being contacted by processes of a microglial cell (green) in CX3CR1*^*+/GFP*^ *mouse electroporated in utero with pCAG-IRES-tdTomato. The analyzed trajectories of microglial processes contacting the neuron are shown on the right panel, warm colors label trajectories of somatic contacts, while cold colors label trajectories of microglial processes contacting neuronal dendrites. The middle panel shows the trajectories overlaid on the recording.*

***Movie S2.*** *Left: stack of electron tomographic 0.5 nm thick virtual sections shows the special nano-architecture of a somatic microglia-neuron junction with closely apposed mitochondria, mitochondria associated membranes (MAMs) and cytoplasmatic structures. The silver-intensified P2Y12R-immunogold grains are clearly visible at the cytoplasmatic surface of microglial membrane. Note that large number of gold particles are clustered exactly where the neuronal cytoplasmatic structure is anchored. Right: 3D model of the same tomographic volume, neuronal membrane is magenta, microglial membrane is green, immunogold particles white, mitochondria light blue, MAM light green, cytoplasmatic densities red, vesicle-like structures blue and microglial reticular membrane structures darker green. Note the specific enrichment of P2Y12R labeling at the core of the junction.*

***Movie S3.*** *Stack of electron tomographic virtual sections shows the special nano-architecture of the core of a somatic microglia-neuron junction. The silver-intensified P2Y12R-immunogold grains are clearly visible at the cytoplasmatic surface of microglial membrane. The tethers between the anchored mitochondria and MAM are clearly visible. The neuronal cytoplasmatic structure (mitochondria, MAM) is anchored to a membrane segment that is precisely facing the high density of P2Y12Rs on the microglial membrane. Note that distance between the neuronal and microglial membrane is the smallest exactly here, and intercellular tethers are also clearly visible.*

***Movie S4.*** *Left: stack of electron tomographic virtual sections shows the special nano-architecture of a somatic microglia-neuron junction with closely apposed mitochondria, MAMs and cytoplasmatic structures. The silver-intensified P2Y12R-immunogold grains are clearly visible at the cytoplasmatic surface of microglial membrane that touches the neuronal cell body, but are present only at a lower density at membrane segments, where the microglia touches a perisomatic bouton. Right: 3D model of the same tomographic volume, neuronal membrane is magenta, microglial membrane is ocker, immunogold particles white, mitochondria light blue, MAM light green, bouton membrane vivid green.*

***Movie S5.*** *In vivo 2P imaging of CX3CR1*^*+/GFP*^ *mice in utero electroporated with CAG-Mito-R-Geco1 construct. Dashed lines show the outlines of two neurons, green microglial processes touch neuronal cell bodies where somatic mitochondria are present. White arrow indicates the contact site of microglia.*

## References

1. M. S. Thion, F. Ginhoux, S. Garel, Microglia and early brain development: An intimate journey. Science. 362, 185–189 (2018).

2. K. Kierdorf, M. Prinz, Microglia in steady state. J. Clin. Invest. 127, 3201–3209 (2017).

3. Y. Zhan et al., Deficient neuron-microglia signaling results in impaired functional brain connectivity and social behavior. Nat. Neurosci. 17, 400–6 (2014).

4. M. W. Salter, B. Stevens, Microglia emerge as central players in brain disease. Nat. Med. 23, 1018–1027 (2017).

5. W. M. Song, M. Colonna, The identity and function of microglia in neurodegeneration. Nat. Immunol. 19, 1048–1058 (2018).

6. D. Davalos et al., ATP mediates rapid microglial response to local brain injury in vivo. Nat. Neurosci. 8, 752–8 (2005).

7. A. Nimmerjahn, Resting Microglial Cells Are Highly Dynamic Surveillants of Brain Parenchyma in Vivo. Science (80-.). 308, 1314–1318 (2005).

8. Y. Wu, L. Dissing-Olesen, B. A. MacVicar, B. Stevens, Microglia: Dynamic Mediators of Synapse Development and Plasticity. Trends Immunol. 36, 605–613 (2015).

9. L. Weinhard et al., Microglia remodel synapses by presynaptic trogocytosis and spine head filopodia induction. Nat. Commun. 9, 1228 (2018).

10. J.-M. Cioni, M. Koppers, C. E. Holt, Molecular control of local translation in axon development and maintenance. Curr. Opin. Neurobiol. 51, 86–94 (2018).

11. T. Misgeld, T. L. Schwarz, Mitostasis in Neurons: Maintaining Mitochondria in an Extended Cellular Architecture. Neuron. 96 (2017), pp. 651–666.

12. M. Terenzio, G. Schiavo, M. Fainzilber, Compartmentalized Signaling in Neurons: From Cell Biology to Neuroscience. Neuron. 96, 667–679 (2017).

13. A. Holtmaat, K. Svoboda, Experience-dependent structural synaptic plasticity in the mammalian brain. Nat. Rev. Neurosci. 10, 759–759 (2009).

14. J. Aarum, K. Sandberg, S. L. B. Haeberlein, M. A. A. Persson, Migration and differentiation of neural precursor cells can be directed by microglia. Proc. Natl. Acad. Sci. 100, 15983–15988 (2003).

15. M. Ueno et al., Layer V cortical neurons require microglial support for survival during postnatal development. Nat. Neurosci. 16, 543–551 (2013).

16. J. L. Marín-Teva, M. A. Cuadros, D. Martín-Oliva, J. Navascués, Microglia and neuronal cell death. Neuron Glia Biol. 7, 25–40 (2011).

17. A. Sierra et al., Surveillance, Phagocytosis, and Inflammation: How Never-Resting Microglia Influence Adult Hippocampal Neurogenesis. Neural Plast. 2014, 1–15 (2014).

18. E. Deutsch et al., Kv2.1 cell surface clusters are insertion platforms for ion channel delivery to the plasma membrane. Mol. Biol. Cell. 23, 2917–2929 (2012).

19. D. P. Mohapatra, K.-S. Park, J. S. Trimmer, Dynamic regulation of the voltage-gated Kv2.1 potassium channel by multisite phosphorylation. Biochem. Soc. Trans. 35, 1064–1068 (2007).

20. A. Menéndez-Méndez et al., Specific Temporal Distribution and Subcellular Localization of a Functional Vesicular Nucleotide Transporter (VNUT) in Cerebellar Granule Neurons. Front. Pharmacol. 8, 951 (2017).

21. R. D. Fields, Nonsynaptic and nonvesicular ATP release from neurons and relevance to neuron-glia signaling. Semin. Cell Dev. Biol. 22, 214–9 (2011).

22. S. E. Haynes et al., The P2Y12 receptor regulates microglial activation by extracellular nucleotides. Nat. Neurosci. 9, 1512–1519 (2006).

23. L. Barna et al., Correlated confocal and super-resolution imaging by VividSTORM. Nat. Protoc. 11, 163–183 (2016).

24. I. D. Campbell, M. J. Humphries, Integrin Structure, Activation, and Interactions. Cold Spring Harb. Perspect. Biol. 3, a004994–a004994 (2011).

25. H. Akiyama, P. L. McGeer, Brain microglia constitutively express beta-2 integrins. J. Neuroimmunol. 30, 81–93 (1990).

26. F. E. McCann et al., The size of the synaptic cleft and distinct distributions of filamentous actin, ezrin, CD43, and CD45 at activating and inhibitory human NK cell immune synapses. J. Immunol. 170, 2862–70 (2003).

27. G. Csordás, D. Weaver, G. Hajnóczky, Endoplasmic Reticulum-Mitochondrial Contactology: Structure and Signaling Functions. Trends Cell Biol. 28, 523–540 (2018).

28. N. S. Chandel, Mitochondria as signaling organelles. BMC Biol. 12, 34 (2014).

29. M. R. Duchen, Mitochondria, calcium-dependent neuronal death and neurodegenerative disease. Pflugers Arch. 464, 111–21 (2012).

30. C. N. Hall, M. C. Klein-Flügge, C. Howarth, D. Attwell, Oxidative phosphorylation, not glycolysis, powers presynaptic and postsynaptic mechanisms underlying brain information processing. J. Neurosci. 32, 8940–51 (2012).

31. U. Ahting et al., The TOM core complex: the general protein import pore of the outer membrane of mitochondria. J. Cell Biol. 147, 959–68 (1999).

32. A. Sugiura, G.-L. McLelland, E. A. Fon, H. M. McBride, A new pathway for mitochondrial quality control: mitochondrial-derived vesicles. EMBO J. 33, 2142–56 (2014).

33. T. Ho et al., Vesicular expression and release of ATP from dopaminergic neurons of the mouse retina and midbrain. Front. Cell. Neurosci. 9, 389 (2015).

34. Y. Moriyama, M. Hiasa, S. Sakamoto, H. Omote, M. Nomura, Vesicular nucleotide transporter (VNUT): appearance of an actress on the stage of purinergic signaling. Purinergic Signal. 13, 387–404 (2017).

35. A. M. Brennan, J. A. Connor, C. W. Shuttleworth, NAD(P)H Fluorescence Transients after Synaptic Activity in Brain Slices: Predominant Role of Mitochondrial Function. J. Cereb. Blood Flow Metab. 26, 1389–1406 (2006).

36. C. W. Shuttleworth, A. M. Brennan, J. A. Connor, NAD(P)H fluorescence imaging of postsynaptic neuronal activation in murine hippocampal slices. J. Neurosci. 23, 3196–208 (2003).

37. G. Szalay et al., Microglia protect against brain injury and their selective elimination dysregulates neuronal network activity after stroke. Nat. Commun. 7, 11499 (2016).

38. M. Kislin et al., Reversible Disruption of Neuronal Mitochondria by Ischemic and Traumatic Injury Revealed by Quantitative Two-Photon Imaging in the Neocortex of Anesthetized Mice. J. Neurosci. 37, 333–348 (2017).

39. H. Misonou, Calcium- and Metabolic State-Dependent Modulation of the Voltage-Dependent Kv2.1 Channel Regulates Neuronal Excitability in Response to Ischemia. J. Neurosci. 25, 11184–11193 (2005).

40. B. O’Rourke, Mitochondrial ion channels. Annu. Rev. Physiol. 69, 19–49 (2007).

41. J. O. Onukwufor, D. Stevens, C. Kamunde, Bioenergetic and volume regulatory effects of mitoKATP channel modulators protect against hypoxia-reoxygenation-induced mitochondrial dysfunction. J. Exp. Biol. 219, 2743–51 (2016).

42. Y. Li, X.-F. Du, C.-S. Liu, Z.-L. Wen, J.-L. Du, Reciprocal regulation between resting microglial dynamics and neuronal activity in vivo. Dev. Cell. 23, 1189–202 (2012).

43. R. D. Stowell et al., Cerebellar microglia are dynamically unique and survey Purkinje neurons in vivo. Dev. Neurobiol. 78, 627–644 (2018).

44. T. E. Gunter, L. Buntinas, G. Sparagna, R. Eliseev, K. Gunter, Mitochondrial calcium transport: mechanisms and functions. Cell Calcium. 28, 285–296 (2000).

45. E. I. Rugarli, T. Langer, Mitochondrial quality control: a matter of life and death for neurons. EMBO J. 31, 1336–49 (2012).

46. P. D. Bhola, A. Letai, Mitochondria-Judges and Executioners of Cell Death Sentences. Mol. Cell. 61, 695–704 (2016).

47. A. Kasahara, L. Scorrano, Mitochondria: from cell death executioners to regulators of cell differentiation. Trends Cell Biol. 24, 761–70 (2014).

48. G. R. Bantug et al., Mitochondria-Endoplasmic Reticulum Contact Sites Function as Immunometabolic Hubs that Orchestrate the Rapid Recall Response of Memory CD8 + T Cells. Immunity. 48, 542–555.e6 (2018).

49. D. Arnoult, F. Soares, I. Tattoli, S. E. Girardin, Mitochondria in innate immunity. EMBO Rep. 12, 901–910 (2011).

50. A. U. Joshi, D. Mochly-Rosen, Mortal engines: Mitochondrial bioenergetics and dysfunction in neurodegenerative diseases. Pharmacol. Res. 138, 2–15 (2018).

51. J. Rieusset, Mitochondria-associated membranes (MAMs): An emerging platform connecting energy and immune sensing to metabolic flexibility. Biochem. Biophys. Res. Commun. 500, 35–44 (2018).

52. M. Krols et al., Mitochondria-associated membranes as hubs for neurodegeneration. Acta Neuropathol. 131, 505–23 (2016).

53. R. Bravo-Sagua et al., Cell death and survival through the endoplasmic reticulum-mitochondrial axis. Curr. Mol. Med. 13, 317–29 (2013).

54. X. Zhang, Y. Chen, C. Wang, L.-Y. M. Huang, Neuronal somatic ATP release triggers neuron-satellite glial cell communication in dorsal root ganglia. Proc. Natl. Acad. Sci. 104, 9864–9869 (2007).

55. G.-L. McLelland, S. A. Lee, H. M. McBride, E. A. Fon, Syntaxin-17 delivers PINK1/parkin-dependent mitochondrial vesicles to the endolysosomal system. J. Cell Biol. 214, 275–91 (2016).

56. V. J. J. Cadete et al., Formation of mitochondrial-derived vesicles is an active and physiologically relevant mitochondrial quality control process in the cardiac system. J. Physiol. 594, 5343–62 (2016).

57. L. Feinshreiber, D. Singer-Lahat, U. Ashery, I. Lotan, Voltage-gated Potassium Channel as a Facilitator of Exocytosis. Ann. N. Y. Acad. Sci. 1152, 87–92 (2009).

58. P. D. Fox et al., Induction of stable ER-plasma-membrane junctions by Kv2.1 potassium channels. J. Cell Sci. 128, 2096–105 (2015).

59. H. Misonou et al., Regulation of ion channel localization and phosphorylation by neuronal activity. Nat. Neurosci. 7, 711–718 (2004).

60. P. J. Mulholland et al., Glutamate Transporters Regulate Extrasynaptic NMDA Receptor Modulation of Kv2.1 Potassium Channels. J. Neurosci. 28, 8801–8809 (2008).

61. C. Madry et al., Microglial Ramification, Surveillance, and Interleukin-1β Release Are Regulated by the Two-Pore Domain K + Channel THIK-1. Neuron. 97, 299–312.e6 (2018).

62. G. Marchal et al., Prolonged persistence of substantial volumes of potentially viable brain tissue after stroke: a correlative PET-CT study with voxel-based data analysis. Stroke. 27, 599–606 (1996).

63. J. C. Baron, M. E. Moseley, For how long is brain tissue salvageable? Imaging-based evidence. J. Stroke Cerebrovasc. Dis. 9, 15–20 (2000).

64. A. Bunevicius, H. Yuan, W. Lin, The Potential Roles of 18 F-FDG-PET in Management of Acute Stroke Patients. Biomed Res. Int. 2013, 1–14 (2013).

65. C. Iadecola, J. Anrather, The immunology of stroke: From mechanisms to translation. Nat. Med. (2011), doi:10.1038/nm.2399.

66. G. Kato et al., Microglial Contact Prevents Excess Depolarization and Rescues Neurons from Excitotoxicity. eNeuro. 3 (2016), doi:10.1523/ENEURO.0004-16.2016.

67. R. Fekete et al., Microglia control the spread of neurotropic virus infection via P2Y12 signalling and recruit monocytes through P2Y12-independent mechanisms. Acta Neuropathol. 136, 461–482 (2018).

68. L. Shao et al., Mitochondrial involvement in psychiatric disorders. Ann. Med. 40, 281–295 (2008).

69. X. Wang, C. Zhang, G. Szábo, Q.-Q. Sun, Distribution of CaMKIIα expression in the brain in vivo, studied by CaMKIIα-GFP mice. Brain Res. 1518, 9–25 (2013).

70. A. H. Meyer, I. Katona, M. Blatow, A. Rozov, H. Monyer, In vivo labeling of parvalbumin-positive interneurons and analysis of electrical coupling in identified neurons. J. Neurosci. 22, 7055–64 (2002).

71. G. López-Bendito et al., Preferential origin and layer destination of GAD65-GFP cortical interneurons. Cereb. Cortex. 14, 1122–33 (2004).

72. H. Dana et al., Thy1-GCaMP6 Transgenic Mice for Neuronal Population Imaging In Vivo. PLoS One. 9, e108697 (2014).

73. T. Matsuda, C. L. Cepko, Electroporation and RNA interference in the rodent retina in vivo and in vitro. Proc. Natl. Acad. Sci. 101, 16–22 (2004).

74. J. Wu et al., Improved orange and red Ca^2^± indicators and photophysical considerations for optogenetic applications. ACS Chem. Neurosci. 4, 963–72 (2013).

75. A. Denes, N. Humphreys, T. E. Lane, R. Grencis, N. Rothwell, Chronic Systemic Infection Exacerbates Ischemic Brain Damage via a CCL5 (Regulated on Activation, Normal T-Cell Expressed and Secreted)-Mediated Proinflammatory Response in Mice. J. Neurosci. 30, 10086–10095 (2010).

76. K. L. Schaar, M. M. Brenneman, S. I. Savitz, Functional assessments in the rodent stroke model. Exp. Transl. Stroke Med. 2, 13 (2010).

77. J. B. Bederson et al., Rat middle cerebral artery occlusion: evaluation of the model and development of a neurologic examination. Stroke. 17, 472–6 (1986).

78. A. Denes et al., AIM2 and NLRC4 inflammasomes contribute with ASC to acute brain injury independently of NLRP3. Proc. Natl. Acad. Sci. 112, 4050–4055 (2015).

79. B. W. McColl, H. V. Carswell, J. McCulloch, K. Horsburgh, Extension of cerebral hypoperfusion and ischaemic pathology beyond MCA territory after intraluminal filament occlusion in C57Bl/6J mice. Brain Res. (2004), doi:10.1016/j.brainres.2003.10.028.

80. J. R. Kremer, D. N. Mastronarde, J. R. McIntosh, Computer visualization of three-dimensional image data using IMOD. J. Struct. Biol. 116, 71–6 (1996).

81. A. Dani, B. Huang, J. Bergan, C. Dulac, X. Zhuang, Superresolution imaging of chemical synapses in the brain. Neuron. 68, 843–56 (2010).

82. B. Dudok et al., Cell-specific STORM super-resolution imaging reveals nanoscale organization of cannabinoid signaling. Nat. Neurosci. 18, 75–86 (2015).

83. J. V. Cramer et al., In vivo widefield calcium imaging of the mouse cortex for analysis of network connectivity in health and brain disease. bioRxiv, 459941 (2018).

84. M. W. Cole, S. Pathak, W. Schneider, Identifying the brain’s most globally connected regions. Neuroimage. 49, 3132–48 (2010).

85. M. Rubinov, O. Sporns, Weight-conserving characterization of complex functional brain networks. Neuroimage. 56, 2068–79 (2011).

86. D. Hinkle, W. Wiersma, S. Jurs, Applied Statistics for the Behavioural Sciences (2003).

